# A Phenome-wide Mendelian Randomisation study on genetically determined serum urate levels in UK Biobank cohort

**DOI:** 10.1101/630293

**Authors:** Xue Li, Xiangrui Meng, Yazhou He, Athina Spiliopoulou, Maria Timofeeva, Wei-Qi Wei, Aliya Gifford, Tian Yang, Tim Varley, Ioanna Tzoulaki, Peter Joshi, Joshua C. Denny, Paul Mckeigue, Harry Campbell, Evropi Theodoratou

## Abstract

**Introduction:** The role of serum urate level has been extensively investigated in observational studies. However, the extent of any causal effect remains unclear, making it difficult to evaluate its clinical relevance.

**Objectives:** To explore any causal or pleiotropic association between serum urate level and a broad spectrum of disease outcomes.

**Methods:** Phenome-wide association study (PheWAS) together with a Bayesian analysis of tree-structured phenotypic models (TreeWAS) was performed to examine disease outcomes related to genetically determined serum urate levels in 339,256 UK Biobank participants. Mendelian Randomisation (MR) analyses were performed to replicate significant findings using various GWAS consortia data. Sensitivity analyses were conducted to examine possible pleiotropic effects on metabolic traits of the genetic variants used as instruments for serum urate.

**Results:** PheWAS analysis, examining the association with 1,431 disease outcomes, identified a multitude of disease outcomes including gout, hypertension, hypercholesterolemia, and heart diseases (e.g., coronary atherosclerosis, myocardial infarction, and ischaemic heart disease) that were associated (p<3.35e-04) with genetically determined serum urate levels. TreeWAS analysis, examining 10,750 ICD-10 diagnostic terms, identified more sub-phenotypes of cardiovascular and cerebrovascular diseases (e.g., angina pectoris, heart failure, cerebral infarction). MR analysis successfully replicated the association with gout, hypertension, heart diseases and blood lipid levels, but indicated the existence of genetic pleiotropy. Sensitivity analyses support an inference that pleiotropic effects of genetic variants on urate and metabolic traits contribute to the observational associations with cardiovascular diseases.

**Conclusions:** High serum urate levels are associated with different types of cardiac events. The finding of genetic pleiotropy indicates the existence of common upstream pathological elements influencing both urate and metabolic traits, and this may suggest new opportunities and challenges for developing drugs targeting a more distal mediator that would be beneficial for both the treatment of gout and the prevention of cardiovascular comorbidities.

## INTRODUCTION

The role of urate has been explored in a large number of observational studies in relation to a multitude of health outcomes.^1^ Apart from gout, compelling evidence exists for the associations between high serum urate level and an increased risk of non-crystal deposition disorders, including hypertension, cardiovascular diseases, and metabolic syndrome.^2, 3^ Although considerable research efforts have been made in trying to understand the pathological role of urate in such disorders, its causal role has not been clearly established. Therefore, it has been argued that either these associations are confounded by other risk factors, such as obesity, or they represent reverse causality.^4^

As is typical in complex traits, genetic determinants are implicated in the regulation of serum urate levels. Genetic studies among twins and families have reported a substantial heritable component of serum urate level with an estimated heritability of 40-60%.^5, 6^ The genetic determinants of serum urate level have been explored in several genome-wide association studies (GWAS) ^7-10^ and the wealth of resultant data allows for the identification and application of genetic variants as instruments to help separate causal from non-causal associations, given that genotypes are generally independent of environmental exposures and the transmission of genetic information is usually unidirectional. Investigating the associations between genetic variants related to serum urate and disease outcomes might help provide causal evidence in support of the hypotheses which links urate to multiple clinical disorders. Previous MR studies using the genetic variants as instruments of serum urate levels reported inconsistent findings. While some supported a causal effect on health outcomes beyond gout (e.g., diabetic macro-vascular disease, cardiovascular disease mortality, and sudden cardiac death), the majority reported no causal relationships.^1^ The generally neutral results indicate that the large effects have probably not been missed, but some of the MR studies could have been underpowered to detect modest effects or suffered from methodological limitations.

Our recently published MR-PheWAS analysis on the interim release data of UK Biobank (n=120,091) provided an overview of the disease outcomes that were associated with the urate genetic risk loci.^11^ Our study demonstrated that serum urate level shared the same genetic risk loci with multiple disease outcomes, particularly those related to cardiovascular/metabolic diseases and autoimmune disorders.^11^ These findings provide a rationale for further investigating whether these cross-phenotype associations are causal. Although we have applied multiple methodologies to distinguish the PheWAS associations that were causal from those due to pleiotropy or genetic linkage, the use of the interim release data of UK Biobank set power limitations to our investigation and did not allow us to investigate less prevalent phenotypes. The release of the full UK Biobank GWAS genotype dataset provides a unique opportunity to further explore the previous MR-PheWAS findings, repeat analysis with the larger available cohort, and include phenotypes that were not investigated in the previous study due to insufficient number of cases.

In this study, we performed an updated Phenome-wide Mendelian randomisation study (PWMR) by using data from the full UK Biobank cohort. A weighted polygenic risk score (GRS), incorporating effect estimates of multiple genetic risk loci taken from the most recent and largest GWAS of serum urate ^8^ was employed as a proxy of serum urate level. The framework of phenome was defined by using both the PheCODE schema (also used in the previous MR-PheWAS) ^11^ and a novel Bayesian analysis framework, termed TreeWAS (tree-structured phenotypic model).^12^ Any replication of previous findings and/or novel findings were further explored in this study.

## METHODS

### UK Biobank

UK Biobank is a large-scale, population-based prospective cohort study, which recruited over 500,000 participants aged between 40-69 years in 2006-2010 and combined extensive measurement of baseline data and genotype data with linked national medical records (e.g. in-patient hospital episode records, cancer registry and death registry) for longitudinal follow-up. This study was constrained to a subset of unrelated White British individuals with high quality genotype data in order to minimize the influence of diverse population structure within UK Biobank. Details about genotype data and phenotype data and the procedures of quality control are described in the **Supplementary material online, Methods** (page 1).

### Weighted genetic risk score

To generate a genetic proxy for serum urate, genetic variants associated with urate were searched across the GWAS catalogue and literature. Thirty one genetic variants associated with urate among European populations were identified from previous GWASs,^7, 8^ and were selected as components of the genetic proxy for serum urate level. The overall proportion of variance (adjusted R^2^) of urate explained by the 31 genetic variants was around 7%.^8^ The SNP effect on urate (effect size and standard error [SE]) was taken from the largest meta-analysis of GWAS performed by the Global Urate Genetics Consortium (GUGC).^8^ A weighted GRS was constructed by incorporating effect estimates of the 31 urate variants for UK Biobank participants. Specifically, the polygenic risk score was created by adding up the number of urate-increasing alleles for each SNP weighted for the SNP effect size on serum urate level (regression beta coefficients) and then adding this weighted score for all 31 SNPs.

### Phenome framework

We analyzed three phenotypic datasets (i.e. in-patient hospital records, cancer registry data, and death registry data) available in the UK Biobank database. As we were interested in disease phenotypes, the ontology of the phenome was defined based on the ICD codes in the electronic medical records. We pooled the hospital episode data, cancer registry data and death registry data together and included both the primary and secondary ICD codes. Individual ICD codes could not be directly used to define the phenome, as they represent specific sub-phenotypes of a similar set of diseases, instead of independent phenotypes. To account for the correlations between ICD codes, we applied two strategies: (i) the PheCODE schema that has been recently updated and successfully adopted in our previous MR-PheWAS;^11^ and (ii) a novel Bayesian analysis of a tree-structured phenotypic model (TreeWAS) that was developed by researchers from the Wellcome Trust Centre for Human Genetics.^12^

#### PheCODE schema

The PheCODE system was developed to combine one or more related ICD codes into distinct disease groups.^13^ To develop a phenotyping method applicable to the ICD-10 coding system in UK Biobank, we created a map to match ICD-9/10 codes to phecodes.^11^ The latest version of the PheCODE system includes 1,866 hierarchical phenotype codes that could be directly matched to the ICD-9/10 codes and provides a scheme to automatically exclude patients that have similar or potentially overlapping disease states from the corresponding control group (e.g., excluding type 1 diabetes from being in control group when analyzing the phenotype of type 2 diabetes).

#### Tree-structured phenotypic model

A novel Bayesian analysis on a tree-structured phenotypic model has recently been developed to interrogate the increasingly specific sub-phenotypes defined by the ICD-10 coding system. It has been suggested that this model has higher statistical power for detecting genotype-phenotype associations ^12^. In principle, this phenotyping method models the genetic coefficients across all phenotypes as a set of random variables. To model the correlations of the hierarchical tree-like structure of ICD-10 codes (termed as tree-structured phenotypic model), a Markov process is applied to allow the genetic coefficients to evolve down the tree trunk and branches. The tree structure is determined based on the classification hierarchy of the ICD-10 coding system, where each node in the tree represents a clinical term in the classification. More details about the tree-structured phenotyping process are described elsewhere.^12^

### Statistical analysis

To take advantage of both phenotyping models, we explored the association between the weighted GRS of urate and the phenome framework defined by both the PheCODE schema (described as PheWAS analysis) and the tree-structured phenotypic model (described as TreeWAS analysis). The correlation with weighted GRS was examined for a number of potential confounding factors including sex, age, body mass index (BMI), assessment center and the first 5 genetic principal components (PCs). In the PheWAS analysis, associations between weighted GRS and phecodes (with no less than 20 cases) were examined by logistic regression. Given that phenotypes investigated are not totally independent in the PheCODE system, since multiple levels of phenotypic granularity were used for the definition of the case-control groups, we applied the false discovery rate (FDR) method (corresponding to the FDR of q<0.05) to account for multiple comparisons instead of the more stringent Bonfferoni correction.^14^ In the TreeWAS analysis, associations between the weighted GRS and the phenome variables were tested by the Bayesian network analysis at both terminal and internal nodes of the tree structure. The marginal posterior probability (PP) for each node in the tree (where its genetic coefficient was non-zero) and the corresponding maximum posteriori effect estimate with 95% credible interval were determined by using the maximum a posteriori (MAP) estimator. Any association with any node of the tree at the PP≥0.95 was reported for further investigation. Details about the TreeWAS analysis have been described previosuly.^12^ All the statistical analyses were implemented by R 3.3.2.

### Replication in MR-base database

To validate findings, PheWAS associations were further examined in the MR-base database for replication in different populations.^15, 16^ We used this platform to make causal inference by performing two-sample MR analysis using available GWAS consortia data. We applied the simplest MR IVW approach as crude analysis; if there was horizontal pleiotropy that violated the assumptions of the MR IVW, we applied a mixture-of-experts machine learning framework (MR-MoE) to predict the performance of three main classes of MR analytical approaches (mean-based, median-based, and mode-based methods) in the context of different models of pleiotropy, and then selected the most likely unbiased causal estimate for each specific circumstance.^17^ Full details of these MR approaches, including their different assumptions, are provided in **Supplementary material online, Methods** (page 3) **and Table S1**. The schematic presentation of the overall study design is shown in **Figure 1**.

**Figure 1.**
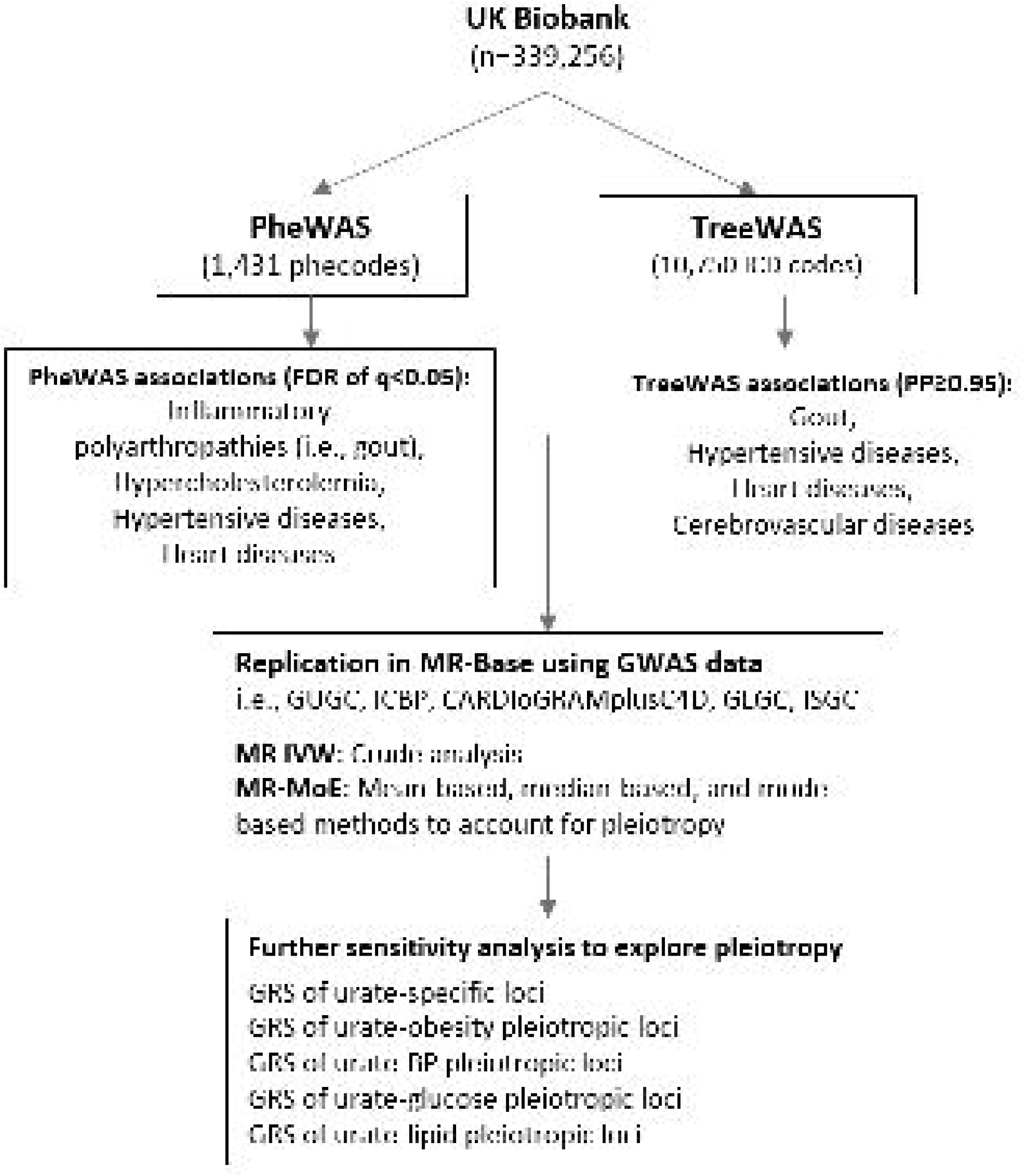
Schematic representation of the study design. (FDR: false discovery rate; PP: posterior probability; GUGC: Global Urate Genetic Consortium; ICBP: International Consortium for Blood Pressure; CARDIoGRAMplusC4D: Coronary ARtery DIsease Genome wide Replication and Meta-analysis [CARDIoGRAM] plus The Coronary Artery Disease [C4D] Genetics consortium; GLGC: Global Lipids Genetic Consortium; ISGC: Ischaemic stroke Genetic Consortium; MR IVW: inverse variance weighted Mendelian randomization; MR-MoE: a mixture-of-experts machine learning framework of Mendelian randomization)

### Sensitivity analysis

We then performed additional sensitivity analyses to further explore any pleiotropic associations. To identify genetic variants showing pleiotropy, we examined their association with a set of metabolic traits (i.e., BMI, waist to hip ratio [WHR], total cholesterol [TC], low-density lipoprotein cholesterol [LDL-c], high-density lipoprotein cholesterol [HDL-c], fasting glucose, 2hr glucose, glycoproteins, systolic blood pressure [SBP], and diastolic blood pressure [DBP]) through publicly available resources from various GWAS consortia (summary of these GWAS are provided in **Supplementary material online, Methods,** page 2). An association was declared as pleiotropic when these GWAS summary data reported any association between the serum urate risk loci and these metabolic traits at p<1.61e-03 (the threshold was determined based on the Bonferroni correction with a significance level of α=0.05 divided by the number of 31 serum urate risk loci analyzed in this study). These 31 urate genetic risk loci were then divided into four categories accordingly: (i) urate-specific loci: including 14 SNPs with no pleiotropic effect on the examined metabolic traits (**Supplementary material online, Table S2**); (ii) urate-obesity pleiotropic loci: including 10 SNPs with pleiotropic effects on BMI or WHR (**Supplementary material online, Table S3**); (iii) urate-BP pleiotropic loci: including 10 SNPs with pleiotropic effects on blood pressures (i.e., DBP and SBP) (**Supplementary material online, Table S4**); (iv) urate-lipid pleiotropic loci: including 6 SNPs with pleiotropic effects on lipids (i.e., TC, LDL-c, HDL-c) (**Supplementary material online, Table S5**); (v) urate-glucose pleiotropic loci: including 3 SNPs with pleiotropic effects on blood glucose (fasting glucose, 2hr glucose, glycoproteins) (**Supplementary material online, Table S6**). A set of GRSs were created accordingly to re-calculate the effect estimates in PheWAS analysis.

## RESULTS

We included 339,256 unrelated White British individuals from the full UK Biobank cohort, consisting of 157,146 men and 182,110 women. The mean age of study population was 56.87 (standard deviation [SD]: 7.99) and the mean BMI was 27.40 (SD: 4.76) kg/m^2^ at the time of recruitment. Other sociodemographic characteristics of the study population are summarized in **Supplementary material online, Table S7**. The mean value of weighted GRS among the study population was 0.44 (SD: 0.31), which is equivalent to 0.44 mg/dL of serum urate level. The correlations between the weighted GRS and potential confounding factors (i.e., age, sex, BMI, assessment center and the PCs) are examined in **Supplementary material online, Table S7**. Of these, two variables (i.e., assessment center and the PCs) were statistically significantly correlated with the weighted GRS and therefore were adjusted as covariates.

### PheWAS and TreeWAS associations

Within the study population, we identified 10,750 unique ICD-10 codes and 3,113 ICD-9 codes in total. After mapping the diagnostic ICD-10/9 codes in UK Biobank to phecodes, the phenome defined by PheCODE schema consisted of 1,853 distinct phecodes among the study population. After filtering out the phecodes with less than 20 cases, PheWAS analysis was performed for 1,431 phecodes (median number of cases: 345 [range: 20-107,298]) which could be classified into 17 broadly related disease categories (**Table 1**). Associations with the weighted GRS of urate were examined for 1,431 case-control groups, leading to an adjusted significance threshold of p<3.35e-04 (corresponding to the FDR of q<0.05) to account for multiple testing. Of these, 13 phecodes were identified to be associated with genetically determined high serum urate level at p<3.35e-04 (**Table 2**). These phecodes represent 4 disease groups: inflammatory polyarthropathies (p=4.97e-19), hypertensive disease (p=6.02e-07), circulatory disease (p=3.29e-04) and metabolic disorders (p= 3.33e-04), and 9 disease outcomes: gout (p=4.27e-123), gouty arthropathy (p=1.39e-05), pyogenic arthritis (p=2.87e-04), essential hypertension (p=6.26e-07), coronary atherosclerosis (p=1.17e-05), ischaemic heart disease (p=1.73e-05), chronic ischaemic heart disease (p=1.52e-05), myocardial infarction (p=5.23e-05), and hypercholesterolemia (p=3.34e-04).

**Table 1.** The number of phenotypes and cases in each disease category.

| Disease categories | Number of phenotypes | Number of cases |  |  |
| --- | --- | --- | --- | --- |
|  |  | Median | Mean | Max. |
| Circulatory diseases | 140 | 434 | 3,581 | 107,298 |
| Congenital anomalies | 45 | 102 | 230 | 1,480 |
| Dermatological diseases | 74 | 283 | 2,544 | 89,976 |
| Diseases in sense organs | 104 | 253 | 1,228 | 31,845 |
| Digestive diseases | 143 | 551 | 3,123 | 62,862 |
| Neoplasms | 129 | 493 | 2,558 | 84,098 |
| Infectious diseases | 48 | 190 | 958 | 8,600 |
| Endocrine and metabolic diseases | 103 | 154 | 1,590 | 35,954 |
| Hematopoietic diseases | 40 | 228 | 1,200 | 10,095 |
| Neurological diseases | 69 | 224 | 1,180 | 32,194 |
| Respiratory diseases | 71 | 674 | 2,448 | 49,782 |
| Mental disorders | 64 | 260 | 1,493 | 23,226 |
| Genitourinary diseases | 140 | 655 | 2,536 | 82,964 |
| Pregnancy complications | 28 | 237 | 914 | 7,518 |
| Musculoskeletal diseases | 109 | 347 | 2,847 | 59,852 |
| Clinical symptoms | 27 | 711 | 3,741 | 33,553 |
| Injuries and poisonings | 97 | 388 | 1,079 | 13,303 |

**Table 2.** Diseases outcomes associated with the weighted GRS of urate in PheWAS analysis.

| **Phecode** | **Disease outcomes** | **n_cases** | **n_controls** | **beta** | **se** | **OR (95%CI)** | **p-value** |
| --- | --- | --- | --- | --- | --- | --- | --- |
| 274.1 | Gout | 2,532 | 335,108 | 1.682 | 0.071 | 5.37 (4.67, 6.18) | 4.27e-123 |
| 714 | Inflammatory polyarthropathies | 15,408 | 320,862 | 0.244 | 0.027 | 1.27 (1.21, 1.34) | 4.97e-19 |
| 401 | Hypertension | 63,694 | 274,477 | 0.076 | 0.015 | 1.07 (1.05, 1.11) | 6.02e-07 |
| 401.1 | Essential hypertension | 63,442 | 274,477 | 0.077 | 0.015 | 1.08 (1.05, 1.11) | 6.26e-07 |
| 411.4 | Coronary atherosclerosis | 25,795 | 311,554 | 0.096 | 0.022 | 1.10 (1.05, 1.14) | 1.17e-05 |
| 274.11 | Gouty arthropathy | 88 | 335,108 | 1.631 | 0.375 | 5.10 (2.45, 10.66) | 1.39e-05 |
| 411.8 | Chronic Ischaemic heart disease | 25,567 | 311,554 | 0.095 | 0.022 | 1.09 (1.05, 1.14) | 1.52e-05 |
| 411 | Ischaemic heart disease | 25,617 | 311,554 | 0.094 | 0.022 | 1.09 (1.05, 1.14) | 1.73e-05 |
| 411.2 | Myocardial infarction | 9,829 | 311,554 | 0.138 | 0.034 | 1.14 (1.07, 1.22) | 5.23e-05 |
| 711.1 | Pyogenic arthritis | 270 | 277,590 | 0.742 | 0.205 | 2.10 (1.41, 3.13) | 2.87e-04 |
| 459.9 | Circulatory disease | 107,298 | 230,622 | 0.046 | 0.013 | 1.04 (1.02, 1.07) | 3.29e-04 |
| 277 | Metabolic disorders | 35,954 | 302,209 | 0.067 | 0.019 | 1.07 (1.03, 1.11) | 3.33e-04 |
| 272.11 | Hypercholesterolemia | 27,040 | 308,948 | 0.077 | 0.021 | 1.08 (1.04, 1.12) | 3.34e-04 |

In the Bayesian analysis framework, containing 10,750 diagnostic terms, a total of 27 parent/child nodes of ICD-10 terms were identified with PP≥0.95. They were clustered mainly in five branches of the hierarchical tree structure (**Figure 2, Supplementary material online, Table S8**): (i) block M10: gout (PP = 1.00) and its sub-phenotypes M10.0 (idiopathic gout) and M10.9 (gout, unspecified); (ii) block I10-I15: hypertensive disease (PP > 0.99) and its sub-phenotype: I10 (essential hypertension); (iii) block I20-I25: ischaemic heart diseases (PP > 0.99) and its sub-phenotypes: I20 (angina pectoris), I21 (acute myocardial infarction), I25 (chronic ischaemic heart disease), I25.1 (atherosclerotic heart disease), I25.2 (old myocardial infarction); (iv) block I30-I52: other forms of heart disease (PP > 0.99) and its sub-phenotype I50 (heart failure) and I50.1 (left ventricular failure); (v) block I60-I69: cerebrovascular diseases (PP > 0.99) and its sub-phenotype I10 (cerebral infarction).

**Figure 2.**
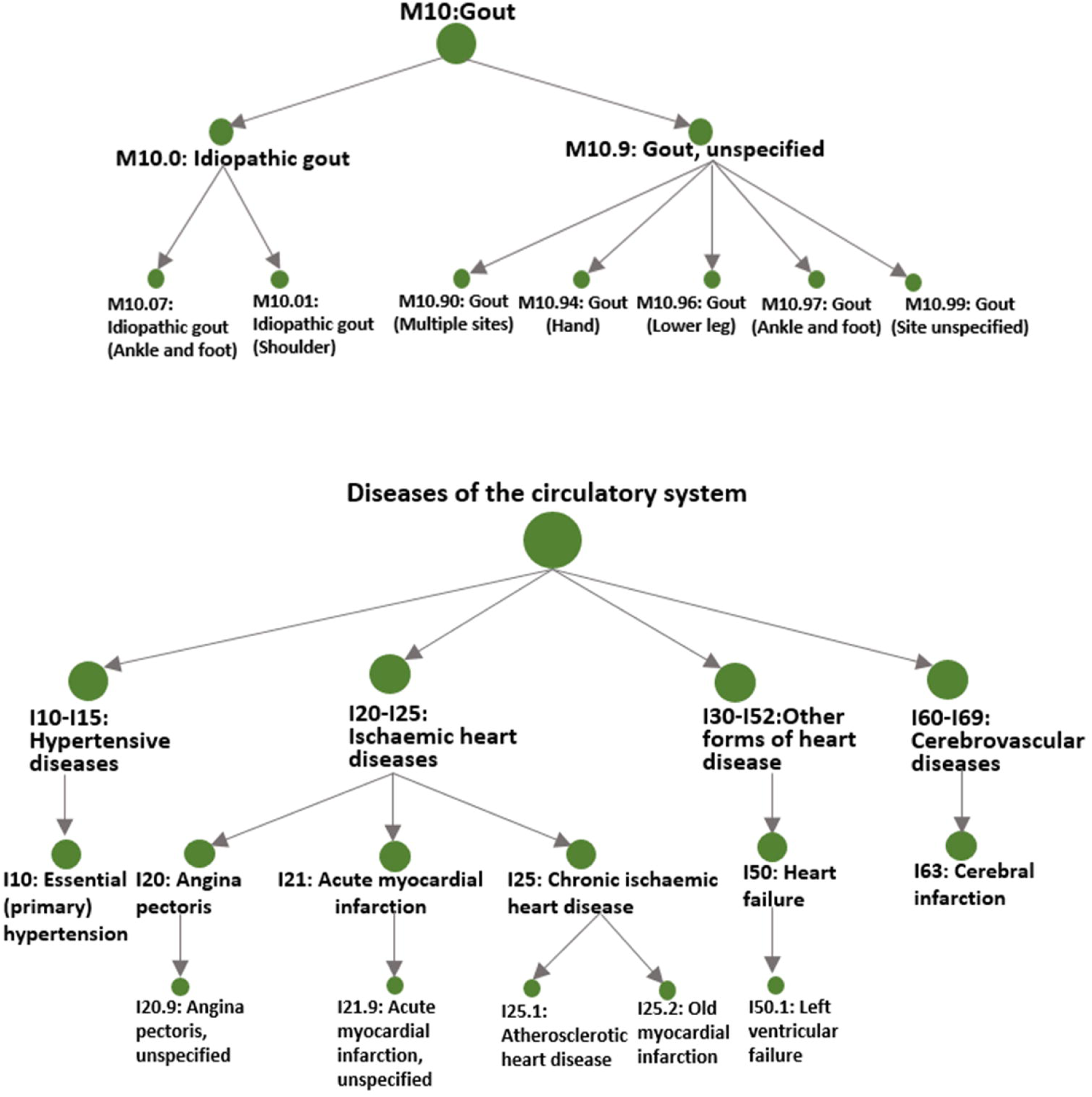
A hierarchical structure of disease outcomes associated with urate in TreeWAS analysis.

Findings from PheWAS and TreeWAS were generally consistent in their associations with gout, hypertensive disease, and heart diseases, while a number of more sub-phenotypes were identified by TreeWAS. Association with the disease group of inflammatory polyarthropathies was statistically significant in PheWAS (OR=1.27, 95%CI: 1.21 to 1.34, p=4.97e-19) but had a moderate PP in TreeWAS (OR=1.07, 95%CI: 1.06 to 1.08, PP=0.76). We examined the specific diseases included in this disease group (M05-M06: rheumatoid arthritis [RA], M07: psoriatic and enteropathic arthropathies, M08-09: juvenile arthritis, M10: gout, and M11-14: arthropathies and other arthritis), and only gout had a significant association in PheWAS analysis. Association with cerebrovascular diseases had a high PP in TreeWAS (OR=1.07, 95%CI: 1.06 to 1.08, PP>0.99) but did not reach the significance threshold of PheWAS (OR=1.08, 95%CI: 0.99 to 1.16, p=0.070), although their estimates were of the same direction. We re-calculated the PheWAS estimates by adding up self-reported stroke cases to increase statistical power, but the corresponding estimates were still not statistically significant (OR=1.05, 95%CI: 0.99 to 1.13, p=0.130).

### Replication in MR-base database

To validate the findings, we performed two-sample MR analyses on associated diseases (i.e., gout, RA, CHD, myocardial infarction, ischaemic stroke) or on their corresponding intermediate traits or surrogate outcomes (i.e., SBP, DBP, TC, LDL-c, HDL-c) (**Table 3**). Results from MR IVW suggested that genetically determined high serum urate level was associated with increased risk of gout (OR=4.53; 95%CI: 3.64 to 5.64; P _*effect*_ = 9.66e-42), DBP (OR=1.04; 95%CI: 1.02 to 1.08; P _*effect*_ =0.044), SBP (OR=1.03; 95%CI: 1.00 to 1.06; P _*effect*_ = 0.050), CHD (OR=1.10; 95%CI: 1.02 to 1.19; *P* _*effect*_ = 0.014), myocardial infarction (OR=1.11; 95%CI: 1.02 to 1.20; P _*effect*_ = 0.017) and decreased level of HDL-c (OR=0.93; 95%CI: 0.88 to 0.98; *P* _*effect*_ = 0.007), but had no effect on ischaemic stroke (OR=1.03; 95%CI: 0.93 to 1.14; *P* _*effect*_ = 0.582). Test for directional horizontal pleiotropy indicated the existence of pleiotropy on the causal estimates of DBP (P _*pleiotropy*_ =0.014), SBP (P _*pleiotropy*_ =0.003), CHD (P _*pleiotropy*_ =0.008), myocardial infarction (P _*pleiotropy*_ =0.008) and HDL-c (P _*pleiotropy*_ =0.016), indicating the MR IVW estimates are likely biased. Using the MR-MOE analysis to select the most appropriate MR approach to deal with different models of pleiotropy, we derived a statistically significant causal estimate only for gout (OR=4.50, 95%CI: 3.62 to 5.59, *P* _*effect*_ =3.35e-77) (Table 3). Causal estimates from each of MR analytical approaches are provided in **Supplementary material online: Table S9-S17**.

**Table 3.** Replication of MR effect estimates in MR-base database.

| Outcome | beta | se | OR (95% CI) | P<sub>effect</sub> | P<sub>pleiotropy</sub>* | n_cases | n_total | Data source |
| --- | --- | --- | --- | --- | --- | --- | --- | --- |
| Replication of significant PheWAS associations |  |  |  |  |  |  |  |  |
| Gout |  |  |  |  |  |  |  |  |
| PheWAS | 1.682 | 0.071 | 5.37 (4.67, 6.18) | 4.27e-123 | -- | 2,532 | 337,640 | UKBB |
| MR IVW | 1.511 | 0.112 | 4.53 (3.64, 5.64) | 9.66e-42 | 0.485 | 2,115 | 67,259 | GUGC |
| MR-MOE | 1.504 | 0.081 | 4.50 (3.62, 5.59) | 3.35E-77 | -- |  |  |  |
| Hypertension |  |  |  |  |  |  |  |  |
| PheWAS | 0.076 | 0.015 | 1.07 (1.05, 1.11) | 6.02e-07 | -- | 63,694 | 338,171 | UKBB |
| DBP |  |  |  |  |  |  |  |  |
| MR IVW | 0.042 | 0.020 | 1.04 (1.02, 1.08) | 0.044 | 0.014 | -- | 69,395 | ICBP |
| MR-MOE | 0.034 | 0.019 | 1.03 (0.99, 1.07) | 0.084 | -- | -- |  |  |
| SBP |  |  |  |  |  |  |  |  |
| MR IVW | 0.031 | 0.015 | 1.03 (1.00, 1.06) | 0.050 | 0.003 | -- | 69,395 | ICBP |
| MR-MOE | 0.008 | 0.007 | 1.01 (0.99, 1.02) | 0.266 | -- | -- |  |  |
| Coronary heart disease (CHD) |  |  |  |  |  |  |  |  |
| PheWAS | 0.094 | 0.022 | 1.09 (1.05, 1.14) | 1.73e-05 | -- | 25,617 | 337,171 | UKBB |
| MR IVW | 0.098 | 0.038 | 1.10 (1.02, 1.19) | 0.014 | 0.008 | 60,801 | 123,504 | CARDIoGRA<br>MplusC4D |
| MR-MOE | 0.047 | 0.028 | 1.05 (0.99, 1.10) | 0.086 | -- |  |  |  |
| Myocardial infarction |  |  |  |  |  |  |  |  |
| PheWAS | 0.138 | 0.034 | 1.14 (1.07, 1.22) | 5.23e-05 | -- | 9,829 | 321,383 | UKBB |
| MR IVW | 0.105 | 0.041 | 1.11 (1.02, 1.20) | 0.017 | 0.008 | 43,676 | 128,199 | CARDIoGRA<br>MplusC4D |
| MR-MOE | 0.058 | 0.030 | 1.06 (0.99, 1.12) | 0.055 | -- |  |  |  |
| Hypercholesterolemia |  |  |  |  |  |  |  |  |
| PheWAS | 0.077 | 0.021 | 1.08 (1.04, 1.12) | 3.34e-04 | -- | 27,040 | 335,988 | UKBB |
| Total cholesterol |  |  |  |  |  |  |  |  |
| MR IVW | 0.028 | 0.036 | 1.03 (0.96, 1.10) | 0.440 | 0.602 | -- | 173,082 | GLGC |
| MR-MOE | 0.005 | 0.026 | 1.00 (0.95, 1.06) | 0.848 | -- | -- |  |  |
| HDL-c |  |  |  |  |  |  |  |  |
| MR IVW | -0.075 | 0.026 | 0.93 (0.88, 0.98) | 0.007 | 0.016 | -- | 187,167 | GLGC |
| MR-MOE | -0.030 | 0.015 | 0.97 (0.94, 1.01) | 0.058 | -- | -- |  |  |
| LDL-c |  |  |  |  |  |  |  |  |
| MR IVW | 0.011 | 0.023 | 1.05 (0.95, 1.16) | 0.627 | 0.175 | -- | 187,365 | GLGC |
| MR-MOE | 0.011 | 0.023 | 1.05 (0.95, 1.16) | 0.627 | -- | -- |  |  |
| Replication of additional TreeWAS associations |  |  |  |  |  |  |  |  |
| Ischaemic stroke |  |  |  |  |  |  |  |  |
| PheWAS | 0.071 | 0.040 | 1.08 (0.99, 1.16) | 0.070 | -- | 9,528 | 338,172 | UKBB |
| MR IVW | 0.029 | 0.052 | 1.03 (0.93, 1.14) | 0.586 | 0.290 | 10,307 | 19,326 | ISGC |
| MR-MOE | 0.029 | 0.052 | 1.03 (0.93, 1.14) | 0.586 | -- |  |  |  |

\* Test for directional horizontal pleiotropy
Abbreviations: UKBB, UK Biobank; GUGC, Global Urate Genetics Consortium; ICBP, International Consortium of Blood Pressure; CARDIoGRAMplusC4D, Coronary ARtery Disease Genome wide Replication and Meta-analysis (CARDIoGRAM) plus The Coronary Artery Disease (C4D) Genetics consortium; GLGC, Global Lipids Genetics Consortium

### Sensitivity analysis

Given that most of the related outcomes were cardiovascular diseases, we performed sensitivity analysis to examine the potential of any pleiotropy effect of urate risk variants on metabolic traits. We re-calculated the PheWAS estimates by using a number of GRSs created based on their association with a set of metabolic traits (**Figure 3, Supplementary material online: Table S18**), and the specific metabolic traits investigated were further determined by the availability of summary GWAS data. The GRS of urate-specific loci was only associated with gout and its upper disease group of inflammatory polyarthropathies, but not with any cardiovascular/metabolic diseases. In contrast, the GRSs of pleiotropic loci on obesity, BP, lipids and glucose showed significant association with both gout and the cardiovascular diseases. Specifically, the GRS of pleiotropic loci on lipids was significantly associated with all cardiovascular diseases, including hypertensive diseases (i.e., essential hypertension), heart diseases (i.e., ischaemic heart diseases), and metabolic disorders (i.e., hypercholesterolemia). Additionally, the GRS of pleiotropic loci on glucose was significantly associated with diabetes (i.e., type 2 diabetes). When removing any group of pleiotropic loci from the creation of GRS, their association with hypertensive diseases, heart diseases, and metabolic disorders were not statistically significant (**Supplementary material online: Table S19**). The effects of pleiotropic loci (mapped with genes) on serum urate level against their effects on four representative disease outcomes were plotted in **Supplementary material online: Figure S1**, in which the two urate transporter genes (*SLC2A9* and *ABCG2*) are recognised as the leading loci driving the association with gout, the *GCKR* gene is the leading locus driving the association with hypercholesterolemia, and the *PTPN11/ATXN2* gene is the leading locus driving the association with hypertension and ischaemic heart diseases.

**Figure 3.**
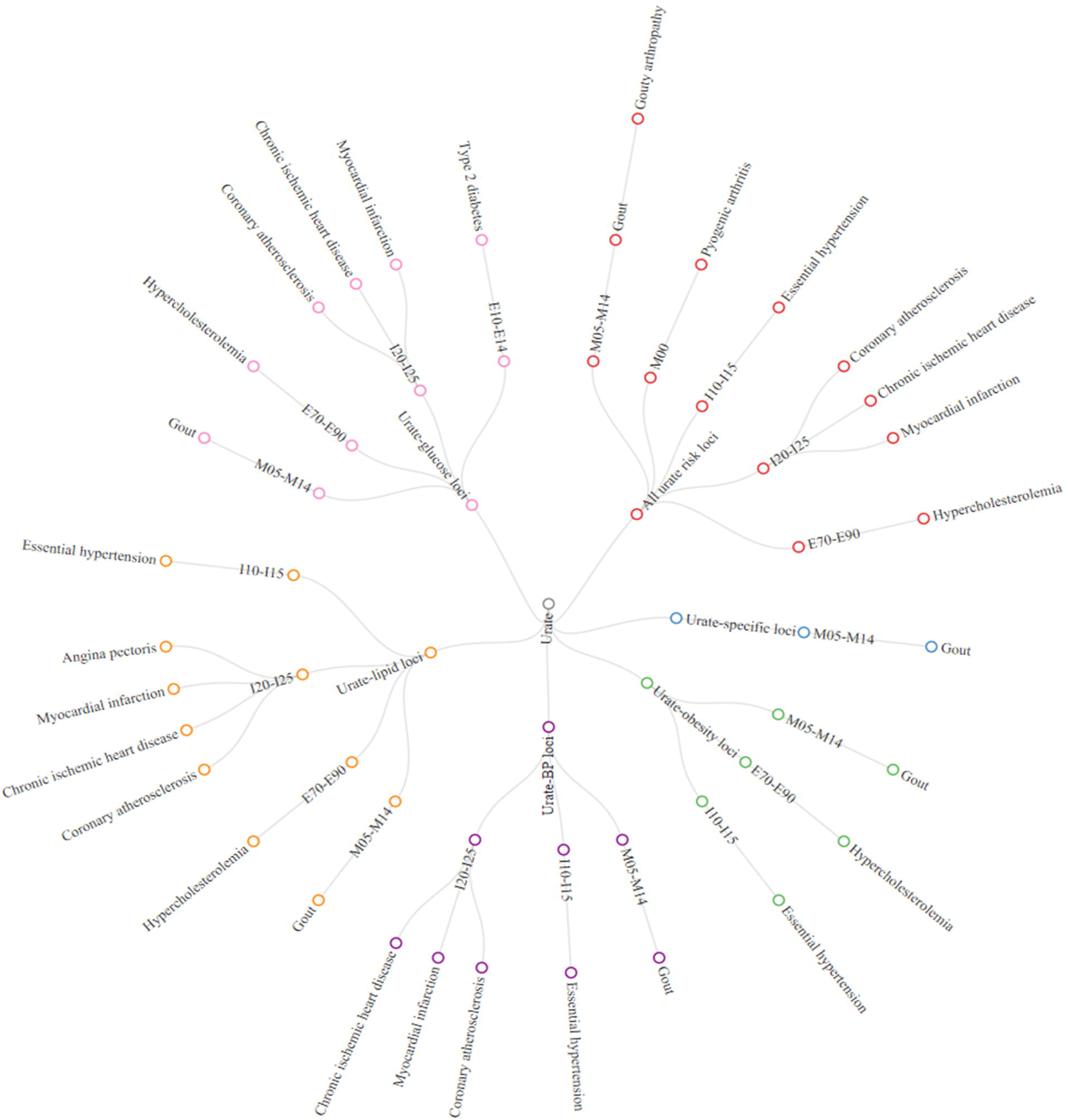
Network plot of the sensitivity analyses of PheWAS using different sets of weighted GRS. The red circles represent the disease outcomes associated with the weighted GRS of the 31 urate genetic rick loci; the blue circles represent disease outcomes associated with the weighted GRS of urate-specific risk loci; the green circles represent diseases outcomes associated with urate-obesity pleiotropic loci; the orange circles represent disease outcomes associated with urate-lipid pleiotropic loci; and the pink circles represent disease outcomes associated with urate-lipid pleiotropic loci. (GRS: genetic risk score; M05-M14: inflammatory polyarthropathies; I10-I15: hypertensive diseases; I20-I25: ischaemic heart diseases; E70-E90: metabolic disorders).

## DISCUSSION

The present study demonstrated that genetically determined high serum urate level was consistently associated with increased risk of several disease groups, including inflammatory polyarthropathies (e.g., gout and gouty arthropathy), hypertensive diseases (e.g., essential hypertension), heart diseases (e.g., coronary atherosclerosis, myocardial infarction, angina pectoris, ischaemic heart disease and heart failure) and metabolic disorders (e.g., hypercholesterolemia). This study using data from the full UK Biobank cohort (n=339,256) verified the associations discovered in the previous MR-PheWAS study based on the interim release of UK Biobank genetic data (n=120,091),^11^ and identified a number of new sub-phenotypes of diseases (e.g., gouty arthropathy, angina pectoris, and heart failure). Some disease outcomes (e.g., disorders of iron metabolism, celiac disease) reported in the previous study were not identified in the present study, as these associations were derived from the genetic linkage disequilibrium between two single variants and therefore were diluted by the use of a weighted GRS of multiple genetic instruments. Association between urate and the risk of gout, hypertension, CHD, myocardial infarction and decreased level of HDL-c were successfully replicated in different European populations by analysing various GWAS consortia data documented in the MR-base database,^15^ but a causal relationship was only supported for gout. Overall, findings from the current study support the observational associations between high serum urate level and increased risk of hypertensive diseases, heart diseases, and metabolic disorders, and also indicated that these associations were more likely due to genetic pleiotropy instead of causality.

A recent umbrella review summarized the published MR studies and examined the causal relationship of serum urate level with a wide range of health outcomes, including gout, cardiovascular, metabolic, and neurocognitive disorders, where for the majority of investigated traits causality was not verified.^1^ There were nine disease outcomes (e.g., diabetic macro-vascular disease, arterial stiffness [internal diameter of carotid artery], adverse renal events, Parkinson’s disease, lifetime anxiety disorders, memory performance, cardiovascular disease mortality, sudden cardiac death, and gout) reported to have a nominally statistically significant causal relationship with urate, but most of them presented with discordant results between MR studies or suffered from methodological limitations (e.g., inadequate study power, invalid genetic instruments) and only that for gout was verified based on convincing evidence.

Specifically, our finding that genetically predicted serum urate level is causally associated with increased risk of gout is not surprising, as it is well known that the causal factor of gout is represented by the monosodium urate crystals (MSU), which leads to acute local inflammation in joints. Moreover, this study also detected an association between urate and the disease group of inflammatory polyarthropathies. To investigate if there was any other type of inflammatory polyarthropathies (beyond gout) associated with urate, we examined the association of urate with all specific diseases included in this group, but none of them were statistically significant. When excluding gout from this disease group, the association was not statistically significant any longer, indicating the observed association was actually driven by gout.

Numerous epidemiological studies have reported that elevated serum urate level is related to increased risk of hypertension and their relationship has been consistent, showing a dose-response relationship and of similar magnitude.^17^ Findings from our current study support this association, but the magnitude of estimated effect size (OR=1.07; 95%CI: 1.05 to 1.11) is smaller than that of traditional epidemiological studies.^18^ In our PheWAS, TreeWAS, and MR IVW analysis we consistently showed a moderate association between urate and different types of heart disease, including coronary atherosclerosis, angina pectoris, ischaemic heart diseases, acute/old myocardial infarction and heart failure, however, the MR-MOE analysis did not support the causal inference after accounting for the presence of pleiotropy.

Large epidemiological studies have established an association between high serum urate level and the increased risk of metabolic disorders.^19^ The NHANES III survey study suggested that high serum urate level was associated with increased level of serum LDL-c, triglycerides, total cholesterol, apolipoprotein-B, and decreased level of HDL-c.^20^ Our study further strengthened this epidemiological evidence and highlighted an association between urate and hypercholesterolemia. Our MR IVW analysis replicated the corresponding association with its surrogate outcome (i.e., HDL-c), but suggested the presence of pleiotropy instead of causality. Additionally, epidemiological studies have also indicated that high serum urate level is associated with increased risk of diabetes.^21^ However, this association was not detected in the main PheWAS or TreeWAS analysis, while sensitivity analysis using the GRS of urate-glucose pleiotropic loci (i.e., *GCKR, IGF1R*, and *SLC16A9*) identified significant association with type 2 diabetes.

To explore how genetic pleiotropy influences the association with cardiovascular/metabolic diseases, we analysed all 31 urate loci across a set of metabolic traits and identified 14 SNPs (urate-specific loci) that were exclusively associated with urate and 17 SNPs (pleiotropic loci) that were associated with metabolic traits. When examining the urate-specific loci, their GRS was only associated with gout and its upper disease group of inflammatory polyarthropathies, but not with any cardiovascular or metabolic diseases. In contrast, when categorizing the pleiotropic loci into different groups (e.g., GRS of urate-obesity loci, GRS of urate-BP loci, GRS of urate-lipid loci and GRS of urate-glucose loci), the GRSs of pleiotropic loci showed consistent associations with both gout and the cardiovascular/metabolic diseases. When removing any group of pleiotropic loci from the creation of GRS (e.g., GRS of urate without pleiotropic loci on BP, or GRS of urate without pleiotropic loci on lipids), their association with heart diseases and metabolic disorders was not statistically significant. Based on these findings, our study suggests that the association between urate and cardiovascular diseases is probably due to the pleiotropic effects of genetic variants on urate and metabolic traits.

Examining the associations between individual urate genetic risk loci and the related disease outcomes highlighted two loci, *GCKR* and *PTPN11/ATXN2* that drive their association with hypercholesterolemia, hypertension and ischaemic heart disease. Pathway network analysis of the leading pleotropic genes provides some clues on how genetic pleiotropy contributes to the association between urate and cardiovascular/metabolic disease. Genetic variation in *GCKR* is shown to be associated with concentrations of urate, triglyceride and glucose ^22^. The most plausible explanation for this observation is that *GCKR* affects both serum urate, triglyceride and glucose levels by a common unconfirmed mediator which is proposed to be glucose-6-phosphate.^23^ The *GCKR* controls the hepatic production of glucose-6-phosphate, which is catabolized for triglyceride synthesis via glycolysis, pyruvate, and acetyl coenzyme A, while glucose-6-phosphate is also a precursor of purine (uric acid) metabolism.^23^ Additionally, gene functional annotation of *PTPN11/ATXN2* highlights another subnetwork around haemostasis pathways, including platelet activation, aggregation, and sensitization (activated by LDL-c),^24^ and these may be relevant to the observed association with hypertension and heart diseases, but how this gene influences serum urate level has not yet been clearly demonstrated.

The detection of a multitude of cross-phenotype associations in this study adds our understanding of the extent of shared genetic/biological components between urate and metabolic traits. Further characterizing the associations between urate and disease outcomes as causal or pleiotropic contributes to our knowledge of how the role of urate should be interpreted and used in clinical practice in the management of related disease conditions. Given that the observational associations between urate and cardiometabolic diseases are more likely due to pleiotropy rather than causality, our study supports the notion that urate could be a predictor but probably not be a direct target for the development of compounds that could reduce cardiovascular/metabolic disease risk. The linked biological pathways between urate and metabolic traits indicated that the frequent co-existence of gout with hypertension, cardiovascular diseases and hyperlipidemia is a range of inter-related disease outcomes due to linked pathogenic components, rather than isolated events. This supports the European League against Rheumatism (EULAR) recommendation of systematic screening and assessment of cardiovascular/metabolic comorbidities in gout patients.^25^ The finding of genetic pleiotropy indicates the existence of common upstream pathological elements influencing both urate and metabolic traits, and this may suggest new opportunities and challenges for developing drugs targeting a more distal mediator that would be beneficial for both the treatment of gout and the prevention of cardiovascular/metabolic comorbidities. This study has focused on the detection of cross-phenotype associations and highlighted the importance of pleiotropy in the links of these complex diseases. We have made efforts to try to understand the cross-phenotype association in the context of a pleiotropy model, but functionally characterizing the underlying biological mechanisms remains a challenge in this field and is worthy of further investigation.

The strengths of this study include its potential to examine a broad spectrum of disease outcomes related to urate and to reflect the shared biological relevance among associated phenotypes, given previous MR studies were typically hypothesis-driven and few studies have comprehensively investigated how serum urate level might influence the overall health. Compared to the previous MR-PheWAS,^11^ the present study extends the prior findings by combining genetic risk loci of urate into a weighted GRS, exploring genetic pleiotropy on a set of metabolic traits systematically, investigating more disease outcomes, assessing their associations with >3-fold more cases, examining consistency of findings across two different phenotyping models to reduce the probability of false positive/negative findings due to factors related to the model, and replicating the findings by performing two-sample MR in different populations. Our study demonstrated the performance of two phenotyping models by accounting for the differences in the specificity and granularity of different phenome definitions and by characterizing the phenotypic correlations among different levels of ICD hierarchy. TreeWAS is shown to increase statistical power and can detect new associations missed by conventional PheWAS.^12^ One of the major accomplishments of this study together with the previous MR-PheWAS have been the establishment of a framework or workflow for PheWAS.^11^ We believe this study would be an excellent starting point for researchers who plan to use the UK Biobank resource to comprehensively interrogate the clinical significance of biomarkers. The updated version of PheCODE schema used in this study is available for researchers who are interested in performing PheWAS in UK Biobank upon request.

This study also has limitations. The causal inference in our study is limited by the common difficulty of pleiotropy caused by the use of multiple genetic instruments. Although we have performed sensitivity analyses by grouping the pleiotropic loci based on metabolic traits and exploring their association separately, there is still a probability of undetected pleiotropy or the possibility that the relatively weak causal effects of urate on diseases were concealed by the strong pleotropic effects of the genetic variants on metabolic traits. Moreover, as most patients (cases) were identified from the inpatient hospital records, this may have impaired the coverage of case ascertainment, especially for the diseases that do not usually cause events for hospitalization. The incorporation of self-reported data would improve this limitation but it is also likely to mistakenly include cases who do not have a true diagnosis and introduce information bias. As UK Biobank is currently performing disease adjudication and processing linkages to general practice records and out-patient data, a widely-covered and accurately-defined criteria of case ascertainment for PheWAS study would be possible in the future.

## Conclusions

Overall, when taken together the findings from PheWAS/TreeWAS, MR replication and sensitivity analyses, we conclude a robust association between urate and a group of diseases, including gout, hypertensive diseases, heart diseases and metabolic disorders of lipids, but the causal role of urate is only supported in gout. Our study indicates that the association between urate and cardiovascular diseases is probably due to the pleiotropic effects of genetic variants on urate and metabolic traits. These findings support that urate could be a good predictor for the cardiovascular/metabolic disease risk. Further investigation on therapies targeting on the shared biological pathways between urate and metabolic traits would be beneficial for the treatment of gout and the primary prevention of cardiovascular/metabolic comorbidities.

## Acknowledgements

The authors would like to thank all participants in the UK Biobank. The authors would also like to thank all the GWAS consortia mentioned in this manuscript for making their data available. The genotype and phenotype data used in this study were obtained from UK Biobank under an approved data request application (application ID: 10775).

## Authors’ contributions

E.T. and H.C. conceived the study and X.L. contributed to the study design. X.L. performed the data analysis. X.L., X.M., W.Q., A.G., J.C.D. and T.V. contributed to the mapping of ICD-10/9 code to phecode. X.L. drafted the manuscript. All authors critically reviewed the manuscript and contributed important intellectual content. All authors have read and approved the final manuscript as submitted.

