## Supplementary material for "A Phenome-wide Mendelian Randomisation study on genetically determined serum urate levels in UK Biobank cohort": Supp

**Supplementary material online**

**Supplementary Methods**

**UK Biobank data**

***Genotype data*** - Genotyping, quality control and genotype imputation were conducted by the UK Biobank team prior to the data release and the exact procedure is described by Bycroft *et al.*^1^ The initial 50,000 participants were genotyped by the Affymetrix UK BiLEVE Axiom array and the remaining 450,000 participants were genotyped by the Affymetrix UK Biobank Axiom array. Genotype imputation was performed based on a merged reference panel of the Haplotype Reference Consortium (HRC)^2^ and the UK10K haplotype resources,^3^ and the classical allelic variations at the MHC region were further imputed by using an additional multi-population reference panel.^4^ For quality control, a list of field variables was made available by the UK Biobank to indicate the genotype quality, population structure, and genetic relatedness.

***Phenotype data*** - A variety of national health systems and sources were used by the UK Biobank to follow up the disease diagnosis, cancer occurrence, and causes of death among the enrolled participants. Currently, there are three main different types of health records (i.e. hospital inpatient episodes, cancer registry data and death registry data) that have been incorporated into the central database. The coding for clinical diagnoses in these datasets followed the World Health Organization’s International Classification of Diseases (ICD) coding systems but used different ICD versions (ICD-10 or ICD-9) according to the date of the record. Primary and/or secondary ICD codes are available in the hospital inpatient data and/or death registry data to classify the main causes and contributory causes of the event of hospitalization and/or death respectively.

**Study population and quality control**

In order to minimize the influence of the diverse population structure in UK Biobank, our study was constrained to a subset of unrelated White British subjects with high quality genotype data. The metrics used for genotype quality control (QC) were based on the data fields created by the UK Biobank. Samples that were identified as a sex mismatch, outliers with high heterozygosity or with high missing rate, putative aneuploidy in sex chromosome, individuals with excess relatives, or non-White British ancestry were all excluded from the analysis. The largest possible subset of individuals without relatedness were identified using an algorithm implemented in the R package “*i-graph (v1.0.1)*” developed by Bycroft and colleagues.^1^

**Other data sources**

**GUGC:** Data on the genetic associations with serum urate levels were obtained from the Global Urate Genetic Consortium (GUGC).^5^ In brief, the GUGC performed meta-analysis of 48 genome-wide association studies (GWAS), totalling 110,347 individuals to assess common variants associated with serum urate levels in European origin. Genotyping was performed on genome-wide chips, and imputation was conducted using HapMap 2 data as the reference.

**GLGC:** Data on the genetic associations with plasma lipid levels were obtained from the Global Lipids Genetic Consortium (GLGC). ^6^ In brief, the GLGC performed a meta-analysis of 46 lipid GWAS and examined subjects of European ancestry, including 94,595 individuals from 23 studies genotyped with GWAS arrays and 93,982 individuals from 37 studies genotyped with the Metabochip array.

**GIANT:** Data on the genetic associations with body mass index (BMI) and waist to hip ratio (WHR) were obtained from the Genetic Investigation of ANthropometric Traits (GIANT) consortium.^7^ In brief, GIANT performed a meta-analysis of 51 GWAS assessing common variants associated with BMI in over 170,000 individuals of European descent. Genotyping was performed using commercially available Affymetrix or Illumina genotyping arrays or custom Perlegen arrays.

**MAGIC:** Data on the genetic associations with plasma glucose was obtained from the Meta-Analyses of Glucose and Insulin-related traits (MAGIC) Consortium.^8^ MAGIC was a genome wide association study (GWAS) that sought to identify genetic determinants of glycemic and metabolic traits. The association between genetic variants and the change in FG (mmol/L) was assessed in 133,010 and 42,854 non-diabetic European individuals^1^. Genotyping was performed using the Metabochip.

**ICBP:** Data on the genetic association with SBP and DBP was obtained from the International Consortium for Blood Pressure (ICBP).^9^ The ICBP-GWAS evaluated associations between 2.5 million genotyped or imputed single nucleotide polymorphisms (SNPs) and SBP and DBP in European ancestry from 29 studies. All studies with GWAS data performed genotyping using commercially available arrays with >300,000 SNPs and were imputed to the HapMap reference panels.

**CARDIoGRAMplusC4D:** Data on the genetic association with the risk of CAD was obtained from the Coronary ARtery DIsease Genome wide Replication and Meta-analysis [CARDIoGRAM] plus The Coronary Artery Disease [C4D] Genetics (CARDIoGRAMplusC4D) Consortium. Briefly, the CARDIoGRAMplusC4D Consortium performed a meta-analysis of 63,746 cases and 130,681 controls^4^. Genotyping was performed using the Metabochip, which is a custom iSELECT chop (Illumina).

**ISGC:** Data on the genetic associations with ischaemic stroke was obtained from the Ischemic stroke Genetic Consortium (ISGC).^10^ They meta-analysed 12 individual genome-wide association studies comprising 10,307 cases and 19,326 controls imputed to the 1000 Genomes (1 KG) phase I reference panel. Genotyping was performed using the Sequenom iPLEX Gold chemistry and genotypes were called using SpectroCHIP array.

**MR-MOE**

Given many MR methods have been developed and each method assumes or performs best for a different model of pleiotropy in order to avoid cherry picking of MR results, we applied a machine learning approach (MR-MoE) to protect MR causal estimates from different patterns of horizontal pleiotropy. Specifically, the MR-MoE considered the 10 MR methods (as described below and in Supplementary Table S1), predicted the performance of each MR method in the context of different models of pleiotropy, and selected the method most likely to be correct for a specific causal analysis. Full details of the MR-MoE are available at <http://mrcieu.github.io/TwoSampleMR/#mr-moe-using-a-mixture-of-experts-machine-learning-approach>.^11^

***Mean-based methods***: Four MR approaches (IVW fixed effects, IVW random effects, Egger fixed effects, Egger random effects) provide four mean-based estimators. The inverse variance weighted (IVW) fixed effects meta-analysis approach assumes that variants exhibit no horizontal pleiotropy. IVW random effects meta-analysis relaxes the horizontal pleiotropy assumption, allowing it to be present but balanced - such that it only leads to increased heterogeneity around the regression line without affecting the slope (and therefore not introducing bias). Fixed effects Egger regression relaxes the horizontal pleiotropy assumption further by allowing a non-zero intercept which essentially allows overall horizontal pleiotropy to be directional, where its total effect influences the outcome in a specific direction.^12^ Random effects Egger regression further allows heterogeneity around the slope having accounted for overall directional horizontal pleiotropy,^13^ as long as the horizontal pleiotropy effects are not correlated with the SNP-exposure effects (also known as the INSIDE assumption).^12^

***Median-based methods***: This analytical approach takes the median effect of all available instruments.^14^ The sample median method requires that half the instruments need to be valid to obtain unbiased estimate. The weighted median method allows stronger instruments to contribute more towards the estimate and obtain an estimates by weighting the contribution of each instrument by the inverse of its variance. The penalised weighted median estimator introduces a further weight to the instruments, penalising any instrument that contributes substantially towards the heterogeneity statistic. Together, this provides three median-based estimators.^14^

***Mode-based methods***: The mode-based estimator clusters the instruments into groups based on similarity of causal effects, and returns the final causal effect estimate based on the cluster that has the largest number of instruments.^15^ This provide three mode-based estimators: the simple mode is the unweighted mode of the empirical density function of causal estimates, the weighted mode is weighted by the inverse variance of the outcome effect, and the penalised weighted mode introduces a further weight to the instruments, penalising any instrument that contributes substantially towards the heterogeneity statistic.

**Supplementary Tables**

Table S1. A summary of MR analytical approaches and their assumptions.

Table S2. A summary of genetic risk variants identified in previous urate GWAS.

Table S3. A summary of pleiotropic loci on urate and obesity traits.

Table S4. A summary of pleiotropic loci on urate and blood pressures (BP).

Table S5. A summary of pleiotropic loci on urate and lipids.

Table S6. A summary of pleiotropic loci on urate and glucose.

Table S7. Association between the GRS of urate and potential confounding factors.

Table S8. Phenotypes associated with the weighted GRS of urate in TreeWAS analysis.

Table S9. Results from MR-MoE analysis for urate and gout

Table S10. Results from MR-MoE analysis for urate and diastolic blood pressure (DBP)

Table S11. Results from MR-MoE analysis for urate and systemic blood pressure (SBP)

Table S12. Results from MR-MoE analysis for urate and coronary heart disease (CHD)

Table S13. Results from MR-MoE analysis for urate and myocardial infarction (MI)

Table S14. Results from MR-MoE analysis for urate and total cholesterol (TC)

Table S15. Results from MR-MoE analysis for urate and high-density lipoprotein cholesterol (HDL_c)

Table S16. Results from MR-MoE analysis for urate and low-density lipoprotein cholesterol (LDL-c)

Table S17. Results from MR-MoE analysis for urate and ischaemic stroke (IS)

Table S18. Sensitivity analysis by including pleiotropic loci on metabolic traits.

Table S19. Sensitivity analysis by excluding the pleiotropic loci of metabolic traits.

**Table S1. A summary of MR analytical approaches and their assumptions**

| **Method type** | **Methods** | **Assumptions** |
| --- | --- | --- |
| Mean-based methods | FE IVW^*^ | No horizontal pleiotropy |
|  | RE IVW^*^ | Balanced horizontal pleiotropy |
|  | FE Egger^*^ | No horizontal pleiotropy after accounting for directional pleiotropy |
|  | RE Egger^*^ | Balanced horizontal pleiotropy after accounting for directional pleiotropy |
|  | Simple mean | Only balanced and/or directional pleiotropy |
| Median-based methods | Simple median | At least half of the instruments are valid. |
|  | Penalised median | At least half the weight of the instruments is due to valid instruments, where each instrument that contributes to high heterogeneity is down weighted. |
|  | Weighted median | At least half the weight of the instruments is due to valid instruments. |
| Mode-based methods | Simple mode | After clustering instruments by causal estimates, the largest cluster is correct, weighting by the exposure variance. |
|  | Weighted mode | After clustering instruments by causal estimates, the largest cluster is correct, weighting by the exposure and outcome variances. |
|  | Penalised mode | After clustering instruments by causal estimates, the largest cluster is correct, where each instrument that contributes to high heterogeneity is down weighted. |

^*^Abbreviations: FE IVW: inverse variance weighted MR with fixed effects; RE IVW: inverse variance weighted MR with random effects; FE Egger: Egger MR with fixed effects; RE Egger: Egger MR with random effects.

**Supplementary Table S2. A summary of genetic risk variants identified in previous urate GWAS.^*^**

| **SNP** | **Chr** | **Closest/GRAIL gene** | **Effect allele** | **Allele freq** | **beta** | **se** | **p-value** | **Pleiotropy** |
| --- | --- | --- | --- | --- | --- | --- | --- | --- |
| rs10821905 | 10 | *A1CF/ASAH2* | A | 0.824 | 0.053 | 0.007 | 3.45E-12 | No |
| rs1165151 | 6 | *SLC17A1/SLC17A3* | T | 0.549 | -0.092 | 0.005 | 4.52E-60 | No |
| rs12498742 | 4 | *SLC2A9/SLC2A9* | A | 0.232 | 0.380 | 0.006 | 0.00E+00 | No |
| rs1394125 | 15 | *UBE2Q2/NRG4* | A | 0.638 | 0.043 | 0.006 | 9.78E-11 | No |
| rs1471633 | 1 | *PDZK1/PDZK1* | A | 0.538 | 0.061 | 0.005 | 1.40E-26 | No |
| rs164009 | 17 | *QRICH2/PRPSAP1* | A | 0.387 | 0.029 | 0.006 | 7.06E-07 | No |
| rs17632159 | 5 | *TMEM171/TMEM171* | C | 0.697 | -0.038 | 0.006 | 2.00E-09 | No |
| rs17786744 | 8 | *STC1/STC1* | A | 0.410 | -0.031 | 0.005 | 8.82E-08 | No |
| rs2078267 | 11 | *SLC22A11/SLC22A11* | T | 0.452 | -0.078 | 0.006 | 8.73E-36 | No |
| rs675209 | 6 | *RREB1/RREB1* | T | 0.731 | 0.063 | 0.006 | 1.38E-21 | No |
| rs6770152 | 3 | *SFMBT1/MUSTN1* | T | 0.424 | -0.048 | 0.006 | 2.66E-16 | No |
| rs7188445 | 16 | *MAF/MAF* | A | 0.672 | -0.032 | 0.006 | 1.15E-07 | No |
| rs7224610 | 17 | *HLF/HLF* | A | 0.396 | -0.038 | 0.006 | 4.74E-11 | No |
| rs742132 | 6 | *LRRC16A/LRRC16A* | A | 0.294 | 0.035 | 0.006 | 1.90E-08 | No |
| rs10480300 | 7 | *PRKAG2/PRKAG2* | T | 0.727 | 0.032 | 0.006 | 9.37E-07 | Yes |
| rs11264341 | 1 | *TRIM46/PKLR* | T | 0.571 | -0.048 | 0.006 | 1.04E-14 | Yes |
| rs1171614 | 10 | *SLC16A9/SLC16A9* | T | 0.769 | -0.074 | 0.007 | 6.48E-23 | Yes |
| rs1178977 | 7 | *BAZ1B/MLXIPL* | A | 0.198 | 0.050 | 0.007 | 6.68E-12 | Yes |
| rs1260326 | 2 | *GCKR/GCKR* | T | 0.607 | 0.077 | 0.006 | 1.31E-40 | Yes |
| rs17050272 | 2 | *INHBB/INHBB* | A | 0.589 | 0.037 | 0.006 | 9.36E-09 | Yes |
| rs2079742 | 17 | *BCAS3/C17orf82* | T | 0.136 | 0.051 | 0.008 | 6.24E-09 | Yes |
| rs2231142 | 4 | *ABCG2/ABCG2* | T | 0.887 | 0.220 | 0.009 | 4.43E-116 | Yes |
| rs2307394 | 2 | *ORC4L/ACVR2A* | T | 0.303 | -0.035 | 0.006 | 7.26E-09 | Yes |
| rs2941484 | 8 | *HNF4G/HNF4G* | T | 0.553 | 0.049 | 0.006 | 3.91E-17 | Yes |
| rs3741414 | 12 | *INHBC/INHBE* | T | 0.755 | -0.071 | 0.007 | 9.79E-22 | Yes |
| rs478607 | 11 | *NRXN2/SLC22A12* | A | 0.153 | -0.048 | 0.007 | 5.31E-10 | Yes |
| rs642803 | 11 | *OVOL1/LTBP3* | T | 0.536 | -0.043 | 0.005 | 4.51E-14 | Yes |
| rs653178 | 12 | *ATXN2/PTPN11* | T | 0.483 | -0.036 | 0.005 | 2.45E-10 | Yes |
| rs6598541 | 15 | *IGF1R/IGF1R* | A | 0.645 | 0.044 | 0.006 | 5.20E-13 | Yes |
| rs7193778 | 16 | *NFAT5/NFAT5* | T | 0.150 | -0.047 | 0.008 | 2.36E-08 | Yes |
| rs729761 | 6 | *VEGFA/VEGFA* | T | 0.715 | -0.046 | 0.006 | 3.05E-12 | Yes |

*GWAS summary data were obtained from the Global Urate Genetic Consortium (GUGC) ^5^; one SNP (rs164009) was included on the basis of its functional role in urate metabolism (encoding a protein involved in the regulation of purine synthesis).

**Supplementary Table S3. A summary of pleiotropic loci on urate and obesity traits.^*^**

| **SNP** | **Chr** | **Closest/GRAIL gene** | **Effect allele** | **BMI** | | | **WHR** | | | **Pleiotropy** |
| --- | --- | --- | --- | --- | --- | --- | --- | --- | --- | --- |
|  |  |  |  | **beta** | **se** | **p-value** | **beta** | **se** | **p-value** |  |
| rs2231142 | 4 | *ABCG2/ABCG2* | T | -0.081 | 0.017 | 2.16E-06 | 0.029 | 0.025 | 0.261 | Yes |
| rs7193778 | 16 | *NFAT5/NFAT5* | T | 0.339 | 0.072 | 2.17E-06 | 0.468 | 0.130 | 3.10E-04 | Yes |
| rs2941484 | 8 | *HNF4G/HNF4G* | T | 0.207 | 0.050 | 3.07E-05 | -0.204 | 0.088 | 0.020 | Yes |
| rs1260326 | 2 | *GCKR/GCKR* | T | -0.131 | 0.032 | 4.22E-05 | 0.130 | 0.045 | 0.004 | Yes |
| rs478607 | 11 | *NRXN2/SLC22A12* | A | -0.281 | 0.069 | 5.08E-05 | 0.152 | 0.119 | 0.200 | Yes |
| rs11264341 | 1 | *TRIM46/PKLR* | T | 0.192 | 0.050 | 1.40E-04 | -0.152 | 0.092 | 0.097 | Yes |
| rs642803 | 11 | *OVOL1/LTBP3* | T | 0.198 | 0.056 | 4.22E-04 | 0.133 | 0.077 | 0.084 | Yes |
| rs653178 | 12 | *ATXN2/PTPN11* | T | -0.233 | 0.067 | 4.93E-04 | 0.072 | 0.097 | 0.458 | Yes |
| rs6598541 | 15 | *IGF1R/IGF1R* | A | 0.194 | 0.057 | 6.89E-04 | 0.089 | 0.077 | 0.251 | Yes |
| rs1178977 | 7 | *BAZ1B/MLXIPL* | A | -0.179 | 0.060 | 0.003 | 0.360 | 0.086 | 2.84E-05 | Yes |
| rs1471633 | 1 | *PDZK1/PDZK1* | A | 0.091 | 0.039 | 0.021 | -0.069 | 0.072 | 0.340 | No |
| rs2079742 | 17 | *BCAS3/C17orf82* | T | -0.158 | 0.069 | 0.022 | -0.039 | 0.129 | 0.762 | No |
| rs3741414 | 12 | *INHBC/INHBE* | T | -0.060 | 0.039 | 0.128 | 0.008 | 0.056 | 0.881 | No |
| rs7224610 | 17 | *HLF/HLF* | A | -0.099 | 0.065 | 0.128 | -0.082 | 0.113 | 0.471 | No |
| rs17050272 | 2 | *INHBB/INHBB* | A | 0.099 | 0.066 | 0.133 | -0.016 | 0.124 | 0.896 | No |
| rs729761 | 6 | *VEGFA/VEGFA* | T | -0.087 | 0.058 | 0.136 | 0.261 | 0.104 | 0.012 | No |
| rs1171614 | 10 | *SLC16A9/SLC16A9* | T | 0.057 | 0.039 | 0.138 | 0.080 | 0.078 | 0.309 | No |
| rs2307394 | 2 | *ORC4L/ACVR2A* | T | 0.106 | 0.075 | 0.159 | -0.031 | 0.129 | 0.807 | No |
| rs1394125 | 15 | *UBE2Q2/NRG4* | A | -0.076 | 0.058 | 0.193 | -0.067 | 0.114 | 0.554 | No |
| rs12498742 | 4 | *SLC2A9/SLC2A9* | A | 0.010 | 0.007 | 0.203 | -0.001 | 0.013 | 0.952 | No |
| rs17786744 | 8 | *STC1/STC1* | A | 0.098 | 0.079 | 0.217 | 0.155 | 0.139 | 0.264 | No |
| rs10480300 | 7 | *PRKAG2/PRKAG2* | T | -0.103 | 0.084 | 0.222 | -0.172 | 0.150 | 0.252 | No |
| rs675209 | 6 | *RREB1/RREB1* | T | -0.046 | 0.043 | 0.280 | -0.206 | 0.079 | 0.009 | No |
| rs742132 | 6 | *LRRC16A/LRRC16A* | A | 0.075 | 0.075 | 0.320 | 0.174 | 0.131 | 0.185 | No |
| rs1165151 | 6 | *SLC17A1/SLC17A3* | T | -0.025 | 0.026 | 0.345 | 0.050 | 0.046 | 0.273 | No |
| rs6770152 | 3 | *SFMBT1/MUSTN1* | T | 0.039 | 0.051 | 0.448 | 0.044 | 0.090 | 0.625 | No |
| rs2078267 | 11 | *SLC22A11/SLC22A11* | T | -0.023 | 0.031 | 0.453 | -0.104 | 0.055 | 0.060 | No |
| rs10821905 | 10 | *A1CF/ASAH2* | A | -0.031 | 0.060 | 0.600 | 0.053 | 0.106 | 0.617 | No |
| rs17632159 | 5 | *TMEM171/TMEM171* | C | 0.027 | 0.069 | 0.694 | -0.016 | 0.126 | 0.901 | No |
| rs7188445 | 16 | *MAF/MAF* | A | 0.011 | 0.080 | 0.893 | -0.203 | 0.141 | 0.149 | No |
| rs164009 | 17 | *QRICH2/PRPSAP1* | A | -0.007 | 0.085 | 0.936 | -0.100 | 0.152 | 0.510 | No |

*GWAS summary data were obtained from the Genetic Investigation of ANthropometric Traits (GIANT) consortium.^7^ Abbreviations: chr, chromosome; BMI, body mass index; WHR, waist to hip ratio.

**Supplementary Table S4. A summary of pleiotropic loci on urate and blood pressure (BP).^*^**

| **SNP** | **Chr** | **Closest/GRAIL gene** | **Effect allele** | **DBP** | | | **SBP** | | | **Pleiotropy** |
| --- | --- | --- | --- | --- | --- | --- | --- | --- | --- | --- |
|  |  |  |  | **beta** | **se** | **p-value** | **beta** | **se** | **p-value** |  |
| rs653178 | 12 | *ATXN2/PTPN11* | T | 1.057 | 0.068 | 6.66E-54 | 0.585 | 0.068 | 1.16E-17 | Yes |
| rs642803 | 11 | *OVOL1/LTBP3* | T | 0.377 | 0.057 | 4.83E-11 | 0.269 | 0.057 | 2.85E-06 | Yes |
| rs2307394 | 2 | *ORC4L/ACVR2A* | T | 0.396 | 0.077 | 2.47E-07 | 0.091 | 0.077 | 0.235 | Yes |
| rs10480300 | 7 | *PRKAG2/PRKAG2* | T | 0.364 | 0.086 | 2.38E-05 | 0.453 | 0.086 | 1.45E-07 | Yes |
| rs729761 | 6 | *VEGFA/VEGFA* | T | 0.236 | 0.060 | 7.38E-05 | -0.040 | 0.060 | 0.506 | Yes |
| rs2941484 | 8 | *HNF4G/HNF4G* | T | 0.194 | 0.051 | 1.30E-04 | 0.155 | 0.051 | 0.002 | Yes |
| rs1178977 | 7 | *BAZ1B/MLXIPL* | A | 0.230 | 0.062 | 1.93E-04 | 0.026 | 0.062 | 0.676 | Yes |
| rs7193778 | 16 | *TRIM46/PKLR* | T | -0.081 | 0.073 | 0.268 | 0.347 | 0.073 | 2.19E-06 | Yes |
| rs11264341 | 1 | *BCAS3/C17orf82* | T | 0.146 | 0.052 | 0.005 | 0.173 | 0.052 | 8.20E-04 | Yes |
| rs2079742 | 17 | *NFAT5/NFAT5* | T | 0.136 | 0.070 | 0.054 | 0.257 | 0.070 | 2.52E-04 | Yes |
| rs6770152 | 3 | *SFMBT1/MUSTN1* | T | 0.154 | 0.052 | 0.003 | 0.119 | 0.052 | 0.022 | No |
| rs7188445 | 16 | *MAF/MAF* | A | -0.214 | 0.082 | 0.009 | -0.037 | 0.082 | 0.655 | No |
| rs7224610 | 17 | *HLF/HLF* | A | 0.174 | 0.067 | 0.009 | 0.202 | 0.067 | 0.002 | No |
| rs12498742 | 4 | *SLC2A9/SLC2A9* | A | 0.017 | 0.008 | 0.023 | 0.009 | 0.008 | 0.219 | No |
| rs1471633 | 1 | *PDZK1/PDZK1* | A | 0.080 | 0.040 | 0.048 | 0.041 | 0.040 | 0.313 | No |
| rs17786744 | 8 | *STC1/STC1* | A | 0.140 | 0.081 | 0.083 | -0.114 | 0.081 | 0.159 | No |
| rs17050272 | 2 | *INHBB/INHBB* | A | 0.111 | 0.068 | 0.099 | 0.063 | 0.068 | 0.354 | No |
| rs3741414 | 12 | *INHBC/INHBE* | T | 0.062 | 0.040 | 0.126 | 0.125 | 0.040 | 0.002 | No |
| rs1165151 | 6 | *SLC17A1/SLC17A3* | T | 0.041 | 0.027 | 0.129 | 0.069 | 0.027 | 0.010 | No |
| rs1171614 | 10 | *SLC16A9/SLC16A9* | T | 0.048 | 0.039 | 0.224 | 0.103 | 0.039 | 0.009 | No |
| rs17632159 | 5 | *TMEM171/TMEM171* | C | -0.063 | 0.071 | 0.368 | -0.066 | 0.070 | 0.349 | No |
| rs10821905 | 10 | *A1CF/ASAH2* | A | 0.046 | 0.061 | 0.447 | 0.174 | 0.061 | 0.004 | No |
| rs478607 | 11 | *NRXN2/SLC22A12* | A | 0.049 | 0.071 | 0.490 | 0.106 | 0.071 | 0.136 | No |
| rs2231142 | 4 | *ABCG2/ABCG2* | T | -0.012 | 0.018 | 0.497 | -0.050 | 0.018 | 0.005 | No |
| rs2078267 | 11 | *SLC22A11/SLC22A11* | T | 0.020 | 0.032 | 0.523 | 0.000 | 0.032 | 0.999 | No |
| rs164009 | 17 | *QRICH2/PRPSAP1* | A | 0.050 | 0.087 | 0.570 | -0.025 | 0.087 | 0.770 | No |
| rs1394125 | 15 | *UBE2Q2/NRG4* | A | 0.026 | 0.060 | 0.658 | 0.042 | 0.060 | 0.477 | No |
| rs6598541 | 15 | *IGF1R/IGF1R* | A | 0.023 | 0.059 | 0.698 | -0.029 | 0.059 | 0.618 | No |
| rs1260326 | 2 | *GCKR/GCKR* | T | -0.012 | 0.033 | 0.714 | 0.066 | 0.033 | 0.044 | No |
| rs675209 | 6 | *RREB1/RREB1* | T | 0.012 | 0.044 | 0.792 | -0.024 | 0.044 | 0.593 | No |
| rs742132 | 6 | *LRRC16A/LRRC16A* | A | 0.005 | 0.077 | 0.950 | 0.003 | 0.077 | 0.966 | No |

*GWAS summary data were obtained from the International Consortium for Blood Pressure (ICBP).^9^

Abbreviations: chr, chromosome; SBP, systolic blood pressure; DBP, diastolic blood pressure.

**Supplementary Table S5. A summary of pleiotropic loci on urate and lipids.^*^**

| **SNPs** | **Chr** | **Closest/GRAIL gene** | **Effect allele** | **TC** | | | **LDL-c** | | | **HDL-c** | | | **Pleiotropy** |
| --- | --- | --- | --- | --- | --- | --- | --- | --- | --- | --- | --- | --- | --- |
|  |  |  |  | **beta** | **se** | **p-value** | **beta** | **se** | **p-value** | **beta** | **se** | **p-value** |  |
| rs1260326 | 2 | *GCKR/GCKR* | T | 0.665 | 0.047 | 6.67E-46 | 0.268 | 0.048 | 2.58E-08 | -0.147 | 0.045 | 1.24E-03 | Yes |
| rs653178 | 12 | *ATXN2/PTPN11* | T | -0.853 | 0.103 | 1.07E-16 | -0.631 | 0.106 | 2.32E-09 | -0.731 | 0.097 | 5.72E-14 | Yes |
| rs17050272 | 2 | *INHBB/INHBB* | A | -0.573 | 0.159 | 3.27E-04 | -0.668 | 0.162 | 3.84E-05 | 0.027 | 0.151 | 0.858 | Yes |
| rs642803 | 11 | *OVOL1/LTBP3* | T | -0.281 | 0.084 | 7.76E-04 | -0.270 | 0.086 | 0.002 | -0.342 | 0.079 | 1.54E-05 | Yes |
| rs3741414 | 12 | *INHBC/INHBE* | T | 0.118 | 0.059 | 0.046 | 0.224 | 0.061 | 2.18E-04 | -0.417 | 0.056 | 1.36E-13 | Yes |
| rs1178977 | 7 | *BAZ1B/MLXIPL* | A | 0.196 | 0.090 | 0.029 | -0.068 | 0.094 | 0.469 | -0.632 | 0.088 | 6.88E-13 | Yes |
| rs6770152 | 3 | *SFMBT1/MUSTN1* | T | -0.302 | 0.108 | 0.005 | -0.244 | 0.110 | 0.027 | -0.142 | 0.102 | 0.165 | No |
| rs7224610 | 17 | *HLF/HLF* | A | -0.287 | 0.137 | 0.036 | -0.258 | 0.139 | 0.064 | -0.105 | 0.129 | 0.414 | No |
| rs729761 | 6 | *VEGFA/VEGFA* | T | 0.243 | 0.128 | 0.058 | 0.254 | 0.133 | 0.055 | -0.317 | 0.122 | 0.009 | No |
| rs17786744 | 8 | *STC1/STC1* | A | 0.300 | 0.168 | 0.074 | 0.319 | 0.171 | 0.062 | -0.213 | 0.158 | 0.178 | No |
| rs1165151 | 6 | *SLC17A1/SLC17A3* | T | -0.087 | 0.055 | 0.117 | -0.055 | 0.057 | 0.327 | -0.124 | 0.052 | 0.018 | No |
| rs2941484 | 8 | *HNF4G/HNF4G* | T | -0.159 | 0.104 | 0.126 | -0.080 | 0.108 | 0.462 | -0.031 | 0.098 | 0.755 | No |
| rs11264341 | 1 | *TRIM46/PKLR* | T | -0.154 | 0.113 | 0.171 | -0.158 | 0.115 | 0.167 | 0.100 | 0.106 | 0.347 | No |
| rs1171614 | 10 | *SLC16A9/SLC16A9* | T | 0.120 | 0.093 | 0.197 | 0.077 | 0.096 | 0.422 | 0.020 | 0.089 | 0.820 | No |
| rs2231142 | 4 | *ABCG2/ABCG2* | T | 0.031 | 0.026 | 0.234 | 0.044 | 0.027 | 0.106 | -0.056 | 0.025 | 0.027 | No |
| rs2079742 | 17 | *BCAS3/C17orf82* | T | 0.163 | 0.149 | 0.275 | 0.086 | 0.153 | 0.573 | 0.214 | 0.141 | 0.130 | No |
| rs1394125 | 15 | *UBE2Q2/NRG4* | A | 0.135 | 0.137 | 0.326 | 0.135 | 0.140 | 0.334 | -0.100 | 0.130 | 0.443 | No |
| rs6598541 | 15 | *IGF1R/IGF1R* | A | -0.070 | 0.084 | 0.402 | -0.050 | 0.089 | 0.573 | -0.248 | 0.082 | 0.002 | No |
| rs478607 | 11 | *NRXN2/SLC22A12* | A | 0.110 | 0.146 | 0.449 | 0.150 | 0.150 | 0.317 | -0.188 | 0.138 | 0.173 | No |
| rs10821905 | 10 | *A1CF/ASAH2* | A | 0.085 | 0.130 | 0.514 | 0.051 | 0.132 | 0.700 | 0.019 | 0.121 | 0.876 | No |
| rs2307394 | 2 | *ORC4L/ACVR2A* | T | -0.094 | 0.151 | 0.534 | -0.071 | 0.154 | 0.643 | -0.083 | 0.143 | 0.562 | No |
| rs164009 | 17 | *QRICH2/PRPSAP1* | A | 0.097 | 0.183 | 0.597 | -0.072 | 0.186 | 0.697 | 0.097 | 0.169 | 0.568 | No |
| rs7193778 | 16 | *NFAT5/NFAT5* | T | 0.081 | 0.157 | 0.608 | -0.028 | 0.160 | 0.862 | -0.253 | 0.147 | 0.085 | No |
| rs7188445 | 16 | *MAF/MAF* | A | -0.078 | 0.175 | 0.655 | -0.019 | 0.178 | 0.916 | 0.091 | 0.163 | 0.577 | No |
| rs675209 | 6 | *RREB1/RREB1* | T | 0.035 | 0.092 | 0.704 | -0.083 | 0.094 | 0.378 | 0.056 | 0.087 | 0.525 | No |
| rs12498742 | 4 | *SLC2A9/SLC2A9* | A | 0.005 | 0.016 | 0.747 | 0.008 | 0.016 | 0.617 | -0.019 | 0.015 | 0.186 | No |
| rs10480300 | 7 | *PRKAG2/PRKAG2* | T | 0.053 | 0.178 | 0.766 | 0.072 | 0.184 | 0.697 | -0.022 | 0.169 | 0.897 | No |
| rs2078267 | 11 | *SLC22A11/SLC22A11* | T | -0.013 | 0.067 | 0.848 | -0.054 | 0.068 | 0.428 | 0.074 | 0.062 | 0.227 | No |
| rs17632159 | 5 | *TMEM171/TMEM171* | C | -0.026 | 0.155 | 0.865 | -0.113 | 0.161 | 0.481 | 0.342 | 0.145 | 0.018 | No |
| rs1471633 | 1 | *PDZK1/PDZK1* | A | -0.008 | 0.089 | 0.926 | 0.054 | 0.090 | 0.549 | -0.190 | 0.082 | 0.020 | No |
| rs742132 | 6 | *LRRC16A/LRRC16A* | A | -0.011 | 0.163 | 0.944 | -0.049 | 0.166 | 0.769 | -0.009 | 0.151 | 0.955 | No |

*GWAS summary data were obtained from the Global Lipids Genetic Consortium (GLGC). ^6^

Abbreviations: chr, chromosome; TC, total cholesterol; LDL-c, low-density lipoprotein cholesterol; HDL-c, high-density lipoprotein cholesterol;

**Supplementary Table S6. A summary of pleiotropic loci on urate and glucose.^*^**

| **SNP** | **Chr** | **Closest/GRAIL gene** | **Effect allele** |  | **Fasting glucose (n=15,234)** | | | **2hr glucose (n=58,074)** | | | **Glycoproteins (n=18,732)** | | **Pleiotropy** |
| --- | --- | --- | --- | --- | --- | --- | --- | --- | --- | --- | --- | --- | --- |
|  |  |  |  | **beta** | **se** | **p-value** | **beta** | **se** | **p-value** | **beta** | **se** | **p-value** |  |
| rs1260326 | 2 | *GCKR/GCKR* | T | 1.182 | 0.247 | 1.67E-06 | -0.416 | 0.040 | 5.57E-25 | -0.078 | 0.151 | 0.603 | Yes |
| rs6598541 | 15 | *IGF1R/IGF1R* | A | -0.682 | 0.455 | 0.134 | 0.273 | 0.075 | 2.77E-04 | 0.297 | 0.251 | 0.236 | Yes |
| rs1171614 | 10 | *SLC16A9/SLC16A9* | T | -0.297 | 0.392 | 0.448 | -0.085 | 0.057 | 0.134 | 0.747 | 0.192 | 1.01E-04 | Yes |
| rs1394125 | 15 | *UBE2Q2/NRG4* | A | -1.302 | 0.512 | 0.011 | 0.009 | 0.084 | 0.912 | -0.302 | 0.295 | 0.306 | No |
| rs17050272 | 2 | *INHBB/INHBB* | A | 1.568 | 0.622 | 0.012 | 0.135 | 0.097 | 0.165 | 0.179 | 0.298 | 0.549 | No |
| rs7224610 | 17 | *HLF/HLF* | A | 1.158 | 0.500 | 0.021 | 0.018 | 0.084 | 0.827 | 0.532 | 0.298 | 0.074 | No |
| rs478607 | 11 | *NRXN2/SLC22A12* | A | -1.083 | 0.521 | 0.038 | 0.025 | 0.088 | 0.775 | 0.189 | 0.276 | 0.493 | No |
| rs653178 | 12 | *ATXN2/PTPN11* | T | 0.833 | 0.528 | 0.114 | -0.053 | 0.089 | 0.553 | -0.044 | 0.302 | 0.883 | No |
| rs3741414 | 12 | *INHBC/INHBE* | T | -0.479 | 0.324 | 0.139 | 0.014 | 0.054 | 0.792 | 0.083 | 0.177 | 0.638 | No |
| rs6770152 | 3 | *SFMBT1/MUSTN1* | T | 0.563 | 0.396 | 0.155 | 0.044 | 0.065 | 0.498 | 0.490 | 0.226 | 0.030 | No |
| rs7188445 | 16 | *MAF/MAF* | A | -0.813 | 0.625 | 0.194 | -0.191 | 0.103 | 0.065 | -0.601 | 0.355 | 0.091 | No |
| rs10480300 | 7 | *PRKAG2/PRKAG2* | T | -0.719 | 0.656 | 0.273 | -0.313 | 0.109 | 0.004 | 0.495 | 0.409 | 0.226 | No |
| rs2231142 | 4 | *ABCG2/ABCG2* | T | -0.150 | 0.145 | 0.302 | 0.031 | 0.024 | 0.199 | 0.104 | 0.093 | 0.264 | No |
| rs1165151 | 6 | *SLC17A1/SLC17A3* | T | 0.207 | 0.207 | 0.317 | 0.068 | 0.034 | 0.042 | 0.112 | 0.116 | 0.335 | No |
| rs164009 | 17 | *QRICH2/PRPSAP1* | A | 0.655 | 0.655 | 0.317 | 0.086 | 0.110 | 0.435 | -0.040 | 0.380 | 0.915 | No |
| rs7193778 | 16 | *NFAT5/NFAT5* | T | 0.447 | 0.574 | 0.437 | 0.136 | 0.096 | 0.155 | 0.050 | 0.324 | 0.878 | No |
| rs1178977 | 7 | *BAZ1B/MLXIPL* | A | -0.340 | 0.480 | 0.479 | -0.122 | 0.080 | 0.127 | -0.268 | 0.279 | 0.338 | No |
| rs2307394 | 2 | *ORC4L/ACVR2A* | T | -0.400 | 0.571 | 0.484 | -0.191 | 0.094 | 0.042 | -0.252 | 0.339 | 0.457 | No |
| rs11264341 | 1 | *TRIM46/PKLR* | T | -0.292 | 0.417 | 0.484 | -0.075 | 0.069 | 0.275 | -0.355 | 0.222 | 0.109 | No |
| rs2078267 | 11 | *SLC22A11/SLC22A11* | T | -0.154 | 0.244 | 0.528 | -0.067 | 0.041 | 0.104 | -0.027 | 0.137 | 0.843 | No |
| rs1471633 | 1 | *PDZK1/PDZK1* | A | -0.197 | 0.311 | 0.528 | 0.036 | 0.051 | 0.478 | -0.035 | 0.173 | 0.839 | No |
| rs742132 | 6 | *LRRC16A/LRRC16A* | A | -0.371 | 0.600 | 0.536 | 0.089 | 0.097 | 0.362 | -0.219 | 0.341 | 0.521 | No |
| rs2941484 | 8 | *HNF4G/HNF4G* | T | 0.180 | 0.388 | 0.643 | 0.067 | 0.063 | 0.287 | -0.021 | 0.219 | 0.924 | No |
| rs675209 | 6 | *RREB1/RREB1* | T | -0.156 | 0.349 | 0.656 | 0.130 | 0.057 | 0.023 | 0.215 | 0.183 | 0.240 | No |
| rs2079742 | 17 | *BCAS3/C17orf82* | T | 0.235 | 0.529 | 0.657 | -0.059 | 0.090 | 0.514 | -0.622 | 0.274 | 0.023 | No |
| rs17786744 | 8 | *STC1/STC1* | A | 0.255 | 0.613 | 0.678 | 0.132 | 0.100 | 0.186 | -0.187 | 0.344 | 0.588 | No |
| rs729761 | 6 | *VEGFA/VEGFA* | T | -0.150 | 0.478 | 0.754 | -0.202 | 0.078 | 0.010 | -0.347 | 0.257 | 0.177 | No |
| rs17632159 | 5 | *TMEM171/TMEM171* | C | -0.161 | 0.579 | 0.782 | -0.050 | 0.092 | 0.587 | -0.093 | 0.317 | 0.769 | No |
| rs10821905 | 10 | *A1CF/ASAH2* | A | 0.045 | 0.453 | 0.920 | 0.079 | 0.075 | 0.294 | -0.413 | 0.241 | 0.087 | No |
| rs642803 | 11 | *OVOL1/LTBP3* | T | -0.014 | 0.442 | 0.975 | 0.014 | 0.072 | 0.847 | 0.068 | 0.249 | 0.785 | No |
| rs12498742 | 4 | *SLC2A9/SLC2A9* | A | -0.001 | 0.058 | 0.985 | 0.000 | 0.009 | 0.978 | 0.029 | 0.033 | 0.376 | No |

*GWAS summary data were obtained from the Meta-Analyses of Glucose and Insulin-related traits (MAGIC) Consortium.^8^

**Supplementary Table S7. Association between the GRS of urate and potential confounding factors.**

| **Continuous variable** | **Mean (SD)** | **Beta (se)** | **p-value** |
| --- | --- | --- | --- |
| Age | 56.87 (7.99) | 0.010 (0.044) | 0.830 |
| BMI | 27.40 (4.76) | -0.023 (0.027) | 0.381 |
| PC1 score | -12.35 (1.61) | 0.007 (0.009) | 0.408 |
| PC2 score | 3.78 (1.50) | -0.023 (0.008) | 0.007 |
| PC3 score | -1.59 (1.58) | -0.003 (0.009) | 0.753 |
| PC4 score | 1.29 (2.94) | 0.104 (0.016) | 1.74e-10 |
| PC5 score | -0.81 (6.61) | 0.344 (0.037) | 2.20e-16 |
| **Categorical variable** | **Levels** | **F-value** | **p-value** |
| Sex | male/female | 0.476 | 0.490 |
| Assessment center | 22 centers | 3.451 | 1.41e-07 |

Abbreviations: BMI, body mass index; PC, (genetic) principal component.

**Supplementary Table S8. Phenotypes associated with the weighted GRS of urate in TreeWAS analysis (PP≥0.95).**

| **ICD-10 coding** | **Disease description** | **max_b^†^** | **b_ci_lhs^†^** | **b_ci_rhs^†^** | **OR (95%CI)** | **PP^*^** |
| --- | --- | --- | --- | --- | --- | --- |
| M10 | M10 Gout | 1.640 | 1.515 | 1.765 | 5.16 (4.55, 5.84) | 1.000 |
| M100 | M10.0 Idiopathic gout | 1.640 | 1.515 | 1.765 | 5.16 (4.55, 5.84) | 0.993 |
| M1007 | M10.07 Idiopathic gout (Ankle and foot) | 1.640 | 1.515 | 1.765 | 5.16 (4.55, 5.84) | 0.993 |
| M109 | M10.9 Gout, unspecified | 1.640 | 1.515 | 1.765 | 5.16 (4.55, 5.84) | 1.000 |
| M1099 | M10.99 Gout, unspecified (Site unspecified) | 1.640 | 1.515 | 1.765 | 5.16 (4.55, 5.84) | 1.000 |
| M1097 | M10.97 Gout, unspecified (Ankle and foot) | 1.640 | 1.515 | 1.765 | 5.16 (4.55, 5.84) | 1.000 |
| M1096 | M10.96 Gout, unspecified (Lower leg) | 1.640 | 1.515 | 1.765 | 5.16 (4.55, 5.84) | 1.000 |
| M1094 | M10.94 Gout, unspecified (Hand) | 1.640 | 1.515 | 1.765 | 5.16 (4.55, 5.84) | 0.993 |
| M1090 | M10.90 Gout, unspecified (Multiple sites) | 1.640 | 1.515 | 1.765 | 5.16 (4.55, 5.84) | 0.985 |
| M1001 | M10.01 Idiopathic gout (Shoulder) | 1.640 | 1.515 | 1.765 | 5.16 (4.55, 5.84) | 1.000 |
| Chapter IX | Chapter IX Diseases of the circulatory system | 0.070 | 0.055 | 0.085 | 1.07 (1.06, 1.09) | 1.000 |
| Block I10-I15 | I10-I15 Hypertensive diseases | 0.070 | 0.055 | 0.085 | 1.07 (1.06, 1.09) | 1.000 |
| I10 | I10 Essential (primary) hypertension | 0.070 | 0.055 | 0.085 | 1.07 (1.06, 1.09) | 1.000 |
| Block I20-I25 | I20-I25 Ischaemic heart diseases | 0.070 | 0.055 | 0.085 | 1.07 (1.06, 1.09) | 1.000 |
| I20 | I20 Angina pectoris | 0.070 | 0.055 | 0.085 | 1.07 (1.06, 1.09) | 0.994 |
| I209 | I20.9 Angina pectoris, unspecified | 0.070 | 0.055 | 0.085 | 1.07 (1.06, 1.09) | 0.972 |
| I21 | I21 Acute myocardial infarction | 0.070 | 0.055 | 0.085 | 1.07 (1.06, 1.09) | 0.994 |
| I219 | I21.9 Acute myocardial infarction, unspecified | 0.070 | 0.055 | 0.085 | 1.07 (1.06, 1.09) | 0.966 |
| I25 | I25 Chronic ischaemic heart disease | 0.070 | 0.055 | 0.085 | 1.07 (1.06, 1.09) | 1.000 |
| I251 | I25.1 Atherosclerotic heart disease | 0.070 | 0.055 | 0.085 | 1.07 (1.06, 1.09) | 0.999 |
| I252 | I25.2 Old myocardial infarction | 0.070 | 0.055 | 0.085 | 1.07 (1.06, 1.09) | 0.987 |
| Block I30-I52 | I30-I52 Other forms of heart disease | 0.070 | 0.055 | 0.085 | 1.07 (1.06, 1.09) | 0.999 |
| I50 | I50 Heart failure | 0.070 | 0.055 | 0.085 | 1.07 (1.06, 1.09) | 0.994 |
| I501 | I50.1 Left ventricular failure | 0.070 | 0.055 | 0.085 | 1.07 (1.06, 1.09) | 0.966 |
| Block I60-I69 | I60-I69 Cerebrovascular diseases | 0.070 | 0.055 | 0.085 | 1.07 (1.06, 1.09) | 0.991 |
| I63 | I63 Cerebral infarction | 0.070 | 0.055 | 0.085 | 1.07 (1.06, 1.09) | 0.951 |

* PP, posterior probability for the beta (β) estimate in the tree analysis not being zero.

† max_b: maximum a posteriori effect estimate (beta) and the 95% credible interval (max_b_lhs, max_b_rhs).

**Table S9. Results from MR-MoE analysis for urate and gout**

| **Method** | **nsnp** | **beta** | **se** | **ci_low** | **ci_upp** | **pval** | **MOE^*^** |
| --- | --- | --- | --- | --- | --- | --- | --- |
| FE IVW | 31 | 1.504 | 0.081 | 1.287 | 1.722 | 3.35E-77 | 0.80 |
| Simple median | 31 | 1.708 | 0.199 | 1.318 | 2.098 | 9.18E-18 | 0.79 |
| RE IVW | 31 | 1.504 | 0.111 | 1.287 | 1.722 | 2.53E-14 | 0.77 |
| Weighted median | 31 | 1.224 | 0.118 | 0.994 | 1.455 | 2.28E-25 | 0.72 |
| Penalised median | 31 | 1.204 | 0.110 | 0.988 | 1.421 | 1.07E-27 | 0.71 |
| FE Egger | 31 | 1.400 | 0.121 | 1.070 | 1.730 | 1.02E-30 | 0.70 |
| Weighted mode | 31 | 1.221 | 0.124 | 0.979 | 1.464 | 6.06E-11 | 0.67 |
| Simple mode | 31 | 1.842 | 0.404 | 1.051 | 2.634 | 7.99E-05 | 0.65 |
| Penalised mode | 31 | 1.221 | 0.116 | 0.994 | 1.449 | 1.37E-11 | 0.56 |
| RE Egger | 31 | 1.400 | 0.168 | 1.070 | 1.730 | 3.61E-09 | 0.42 |

*A predictor for each method for how well it performs in terms of high power and low type 1 error (scaled 0-1, where 1 is best performance) for causal inference.

**Table S10. Results from MR-MoE analysis for urate and diastolic blood pressure (DBP).**

| **Method** | **nsnp** | **beta** | **se** | **ci_low** | **ci_upp** | **pval** | **MOE^*^** |
| --- | --- | --- | --- | --- | --- | --- | --- |
| Simple mode | 31 | 0.034 | 0.019 | -0.003 | 0.070 | 0.084 | 0.88 |
| Weighted mode | 31 | 0.016 | 0.007 | 0.003 | 0.029 | 0.024 | 0.80 |
| Weighted median | 31 | 0.018 | 0.007 | 0.004 | 0.032 | 0.013 | 0.79 |
| Simple median | 31 | 0.050 | 0.019 | 0.013 | 0.086 | 0.007 | 0.78 |
| Penalised median | 31 | 0.018 | 0.007 | 0.003 | 0.032 | 0.015 | 0.76 |
| Penalised mode | 31 | 0.016 | 0.007 | 0.002 | 0.029 | 0.030 | 0.71 |
| FE IVW | 31 | 0.042 | 0.006 | 0.003 | 0.081 | 1.23E-13 | 0.69 |
| FE Egger | 31 | -0.010 | 0.008 | -0.063 | 0.042 | 0.212 | 0.64 |
| RE IVW | 31 | 0.042 | 0.020 | 0.003 | 0.081 | 0.044 | 0.59 |
| RE Egger | 31 | -0.010 | 0.027 | -0.063 | 0.042 | 0.701 | 0.37 |

*A predictor for each method for how well it performs in terms of high power and low type 1 error (scaled 0-1, where 1 is best performance) for causal inference.

**Table S11. Results from MR-MoE analysis for urate and systemic blood pressure (SBP).**

| **Method** | **nsnp** | **beta** | **se** | **ci_low** | **ci_upp** | **pval** | **MOE^*^** |
| --- | --- | --- | --- | --- | --- | --- | --- |
| Weighted mode | 31 | 0.008 | 0.007 | -0.006 | 0.023 | 0.266 | 0.72 |
| Simple mode | 31 | 0.027 | 0.030 | -0.032 | 0.086 | 0.380 | 0.66 |
| Penalised mode | 31 | 0.012 | 0.009 | -0.004 | 0.029 | 0.155 | 0.66 |
| Weighted median | 31 | 0.011 | 0.007 | -0.004 | 0.025 | 0.155 | 0.64 |
| Simple median | 31 | 0.066 | 0.017 | 0.032 | 0.100 | 1.46E-04 | 0.61 |
| Penalised median | 31 | 0.011 | 0.007 | -0.004 | 0.025 | 0.142 | 0.57 |
| FE IVW | 31 | 0.031 | 0.006 | 0.001 | 0.061 | 0.000 | 0.55 |
| RE IVW | 31 | 0.031 | 0.015 | 0.001 | 0.061 | 0.051 | 0.54 |
| FE Egger | 31 | -0.015 | 0.008 | -0.053 | 0.024 | 0.076 | 0.52 |
| RE Egger | 31 | -0.015 | 0.020 | -0.053 | 0.024 | 0.457 | 0.4 |

*A predictor for each method for how well it performs in terms of high power and low type 1 error (scaled 0-1, where 1 is best performance) for causal inference.

**Table S12. Results from MR-MoE analysis for urate and coronary heart disease (CHD).**

| **Method** | **nsnp** | **beta** | **se** | **ci_low** | **ci_upp** | **pval** | **MOE^*^** |
| --- | --- | --- | --- | --- | --- | --- | --- |
| Weighted median | 31 | 0.047 | 0.028 | -0.007 | 0.102 | 0.086 | 0.81 |
| Simple median | 31 | 0.171 | 0.055 | 0.062 | 0.280 | 0.002 | 0.78 |
| Simple mode | 31 | 0.172 | 0.073 | 0.028 | 0.315 | 0.026 | 0.77 |
| Penalised median | 31 | 0.046 | 0.029 | -0.010 | 0.102 | 0.105 | 0.76 |
| FE IVW | 31 | 0.098 | 0.022 | 0.024 | 0.172 | 7.34E-06 | 0.71 |
| Weighted mode | 31 | 0.048 | 0.025 | -0.001 | 0.097 | 0.065 | 0.71 |
| Penalised mode | 31 | 0.048 | 0.026 | -0.004 | 0.100 | 0.080 | 0.71 |
| RE IVW | 31 | 0.098 | 0.038 | 0.024 | 0.172 | 0.014 | 0.68 |
| FE Egger | 31 | -0.002 | 0.032 | -0.099 | 0.094 | 0.939 | 0.47 |
| RE Egger | 31 | -0.002 | 0.049 | -0.099 | 0.094 | 0.961 | 0.35 |

*A predictor for each method for how well it performs in terms of high power and low type 1 error (scaled 0-1, where 1 is best performance) for causal inference.

**Table S13. Results from MR-MoE analysis for urate and myocardial infarction (MI).**

| **Method** | **nsnp** | **beta** | **se** | **ci_low** | **ci_upp** | **pval** | **MOE^*^** |
| --- | --- | --- | --- | --- | --- | --- | --- |
| Weighted median | 31 | 0.058 | 0.030 | -0.001 | 0.117 | 0.055 | 0.81 |
| Simple mode | 31 | 0.215 | 0.083 | 0.053 | 0.377 | 0.014 | 0.79 |
| FE IVW | 31 | 0.105 | 0.024 | 0.024 | 0.186 | 1.45E-05 | 0.78 |
| Simple median | 31 | 0.192 | 0.057 | 0.080 | 0.304 | 0.001 | 0.76 |
| Penalised mode | 31 | 0.047 | 0.030 | -0.012 | 0.106 | 0.125 | 0.73 |
| Penalised median | 31 | 0.056 | 0.030 | -0.003 | 0.115 | 0.064 | 0.72 |
| Weighted mode | 31 | 0.047 | 0.029 | -0.009 | 0.104 | 0.112 | 0.72 |
| RE IVW | 31 | 0.105 | 0.041 | 0.024 | 0.186 | 0.017 | 0.65 |
| FE Egger | 31 | -0.001 | 0.035 | -0.108 | 0.106 | 0.978 | 0.46 |
| RE Egger | 31 | -0.001 | 0.055 | -0.108 | 0.106 | 0.986 | 0.40 |

*A predictor for each method for how well it performs in terms of high power and low type 1 error (scaled 0-1, where 1 is best performance) for causal inference.

**Table S14. Results from MR-MoE analysis for urate and total cholesterol (TC).**

| **Method** | **nsnp** | **beta** | **se** | **ci_low** | **ci_upp** | **pval** | **MOE^*^** |
| --- | --- | --- | --- | --- | --- | --- | --- |
| Simple median | 31 | 0.005 | 0.026 | -0.046 | 0.056 | 0.848 | 0.87 |
| RE IVW | 31 | 0.028 | 0.036 | -0.042 | 0.098 | 0.440 | 0.85 |
| RE Egger | 31 | 0.050 | 0.053 | -0.053 | 0.154 | 0.348 | 0.82 |
| Weighted median | 31 | 0.011 | 0.014 | -0.017 | 0.038 | 0.452 | 0.81 |
| Penalised mode | 31 | 0.011 | 0.013 | -0.016 | 0.037 | 0.433 | 0.80 |
| Simple mode | 31 | 0.045 | 0.043 | -0.039 | 0.129 | 0.302 | 0.79 |
| Penalised median | 31 | 0.011 | 0.014 | -0.017 | 0.039 | 0.442 | 0.77 |
| Weighted mode | 31 | 0.011 | 0.012 | -0.014 | 0.035 | 0.398 | 0.76 |
| FE Egger | 31 | 0.050 | 0.016 | -0.053 | 0.154 | 0.001 | 0.61 |
| FE IVW | 31 | 0.028 | 0.011 | -0.042 | 0.098 | 0.009 | 0.52 |

*A predictor for each method for how well it performs in terms of high power and low type 1 error (scaled 0-1, where 1 is best performance) for causal inference.

**Table S15. Results from MR-MoE analysis for urate and high-density lipoprotein cholesterol (HDL_c).**

| **Method** | **nsnp** | **beta** | **se** | **ci_low** | **ci_upp** | **pval** | **MOE^*^** |
| --- | --- | --- | --- | --- | --- | --- | --- |
| Penalised mode | 31 | -0.030 | 0.015 | -0.059 | 0.001 | 0.058 | 0.74 |
| Weighted mode | 31 | -0.030 | 0.014 | -0.056 | -0.003 | 0.038 | 0.72 |
| RE IVW | 31 | -0.075 | 0.026 | -0.125 | -0.024 | 0.007 | 0.70 |
| RE Egger | 31 | -0.035 | 0.037 | -0.108 | 0.039 | 0.361 | 0.69 |
| Penalised median | 31 | -0.020 | 0.014 | -0.048 | 0.007 | 0.150 | 0.69 |
| Simple mode | 31 | -0.021 | 0.048 | -0.115 | 0.073 | 0.669 | 0.69 |
| Weighted median | 31 | -0.021 | 0.015 | -0.050 | 0.009 | 0.166 | 0.60 |
| Simple median | 31 | -0.083 | 0.032 | -0.145 | -0.021 | 0.009 | 0.48 |
| FE Egger | 31 | -0.035 | 0.015 | -0.108 | 0.039 | 0.021 | 0.43 |
| FE IVW | 31 | -0.075 | 0.010 | -0.125 | -0.024 | 4.09E-13 | 0.42 |

*A predictor for each method for how well it performs in terms of high power and low type 1 error (scaled 0-1, where 1 is best performance) for causal inference.

**Table S16. Results from MR-MoE analysis for urate and low-density lipoprotein cholesterol (LDL-c).**

| **Method** | **nsnp** | **beta** | **se** | **ci_low** | **ci_upp** | **pval** | **MOE^*^** |
| --- | --- | --- | --- | --- | --- | --- | --- |
| Weighted median | 31 | 0.014 | 0.015 | -0.014 | 0.043 | 0.335 | 0.85 |
| RE IVW | 31 | 0.011 | 0.023 | -0.034 | 0.057 | 0.627 | 0.84 |
| Simple median | 31 | -0.049 | 0.029 | -0.105 | 0.007 | 0.089 | 0.78 |
| Weighted mode | 31 | 0.011 | 0.014 | -0.016 | 0.037 | 0.428 | 0.78 |
| Penalised mode | 31 | 0.011 | 0.015 | -0.018 | 0.039 | 0.460 | 0.73 |
| Penalised median | 31 | 0.012 | 0.015 | -0.018 | 0.041 | 0.438 | 0.72 |
| Simple mode | 31 | -0.038 | 0.043 | -0.122 | 0.046 | 0.377 | 0.72 |
| RE Egger | 31 | 0.041 | 0.034 | -0.025 | 0.107 | 0.230 | 0.71 |
| FE IVW | 31 | 0.011 | 0.011 | -0.034 | 0.057 | 0.298 | 0.70 |
| FE Egger | 31 | 0.041 | 0.016 | -0.025 | 0.107 | 0.010 | 0.53 |

*A predictor for each method for how well it performs in terms of high power and low type 1 error (scaled 0-1, where 1 is best performance) for causal inference.

**Table S17. Results from MR-MoE analysis for urate and ischaemic stroke (IS).**

| **Method** | **nsnp** | **beta** | **se** | **ci_low** | **ci_upp** | **pval** | **MOE^*^** |
| --- | --- | --- | --- | --- | --- | --- | --- |
| RE IVW | 31 | 0.029 | 0.052 | -0.073 | 0.131 | 0.586 | 0.81 |
| Simple median | 31 | 0.021 | 0.083 | -0.141 | 0.183 | 0.801 | 0.81 |
| Simple mean | 31 | 0.021 | 0.045 | -0.068 | 0.110 | 0.644 | 0.80 |
| Weighted mode | 31 | -0.018 | 0.047 | -0.110 | 0.074 | 0.699 | 0.74 |
| Weighted median | 31 | -0.011 | 0.049 | -0.107 | 0.086 | 0.829 | 0.72 |
| Penalised median | 31 | -0.012 | 0.046 | -0.103 | 0.079 | 0.796 | 0.71 |
| FE IVW | 31 | 0.029 | 0.038 | -0.073 | 0.131 | 0.447 | 0.65 |
| Penalised mode | 31 | -0.018 | 0.048 | -0.113 | 0.076 | 0.706 | 0.65 |
| Simple mode | 31 | -0.004 | 0.131 | -0.260 | 0.252 | 0.975 | 0.61 |
| FE Egger | 31 | -0.024 | 0.055 | -0.174 | 0.126 | 0.665 | 0.60 |
| RE Egger | 31 | -0.024 | 0.076 | -0.174 | 0.126 | 0.757 | 0.60 |

*A predictor for each method for how well it performs in terms of high power and low type 1 error (scaled 0-1, where 1 is best performance) for causal inference.

**Supplementary Table S18. Sensitivity analysis by including pleiotropic loci on metabolic traits.**

| **Disease outcomes** | **GRS of all-urate loci (n=31)** | | | **GRS of urate-specific loci (n=14)** | | | **GRS of urate-obesity pleiotropic loci (n=10)** | | | **GRS of urate-BP pleiotropic loci (n=10)** | | | **GRS of urate-lipid pleiotropic loci (GRS=6)** | | | **GRS of urate-glucose pleiotropic loci (GRS=3)** | | |
| --- | --- | --- | --- | --- | --- | --- | --- | --- | --- | --- | --- | --- | --- | --- | --- | --- | --- | --- |
|  | **OR (95%CI)** | **p-value** | **FDR** | **OR (95%CI)** | **p-value** | **FDR** | **OR (95%CI)** | **p-value** | **FDR** | **OR (95%CI)** | **p-value** | **FDR** | **OR (95%CI)** | **p-value** | **FDR** | **OR (95%CI)** | **p-value** | **FDR** |
| Gout | 5.37 (4.67, 6.18) | 4.27E-123 | TRUE | 3.77 (3.19, 4.46) | 4.42E-54 | TRUE | 12.82 (9.91, 16.59) | 5.10E-84 | TRUE | 5.15 (3.26, 8.15) | 2.32E-12 | TRUE | 9.52 (6.05, 14.99) | 2.19E-22 | TRUE | 10.83 (6.37, 18.41) | 1.38E-18 | TRUE |
| Inflammatory polyarthropathies | 1.27 (1.21, 1.34) | 4.97E-19 | TRUE | 1.22 (1.15, 1.30) | 6.45E-10 | TRUE | 1.57 (1.40, 1.76) | 3.39E-14 | TRUE | 1.52 (1.26, 1.84) | 1.27E-05 | TRUE | 1.52 (1.26, 1.83) | 1.25E-05 | TRUE | 1.33 (1.07, 1.66) | 0.010 | FALSE |
| Hypertension | 1.07 (1.05, 1.11) | 6.02E-07 | TRUE | 1.03 (1.00, 1.07) | 0.075 | FALSE | 1.14 (1.06, 1.22) | 1.42E-04 | TRUE | 1.72 (1.55, 1.92) | 2.13E-23 | TRUE | 1.45 (1.31, 1.61) | 3.99E-12 | TRUE | 1.10 (0.97, 1.24) | 0.138 | FALSE |
| Essential hypertension | 1.08 (1.05, 1.11) | 6.26E-07 | TRUE | 1.03 (1.00, 1.07) | 0.074 | FALSE | 1.14 (1.07, 1.22) | 1.37E-04 | TRUE | 1.72 (1.55, 1.91) | 2.87E-23 | TRUE | 1.45 (1.31, 1.61) | 4.08E-12 | TRUE | 1.10 (0.97, 1.24) | 0.146 | FALSE |
| Coronary atherosclerosis | 1.10 (1.05, 1.14) | 1.17E-05 | TRUE | 1.05 (1.00, 1.11) | 0.052 | FALSE | 1.18 (1.07, 1.30) | 5.96E-04 | FALSE | 1.38 (1.18, 1.61) | 3.37E-05 | TRUE | 1.80 (1.55, 2.09) | 1.35E-14 | TRUE | 1.45 (1.21, 1.72) | 3.89E-05 | TRUE |
| Gouty arthropathy | 5.10 (2.45, 10.66) | 1.39E-05 | TRUE | 4.38 (1.77, 10.82) | 0.001 | FALSE | 9.83 (2.50, 38.67) | 1.08E-03 | FALSE | 1.94 (0.17, 21.86) | 0.592 | FALSE | 3.28 (0.30, 36.00) | 0.331 | FALSE | 12.57 (0.75, 209.57) | 0.078 | FALSE |
| Chronic Ischaemic heart disease | 1.09 (1.05, 1.14) | 1.52E-05 | TRUE | 1.05 (1.00, 1.10) | 0.057 | FALSE | 1.18 (1.07, 1.30) | 5.79E-04 | FALSE | 1.37 (1.18, 1.59) | 5.49E-05 | TRUE | 1.79 (1.54, 2.09) | 2.64E-14 | TRUE | 1.44 (1.20, 1.71) | 5.81E-05 | TRUE |
| Ischaemic Heart Disease | 1.09 (1.05, 1.14) | 1.73E-05 | TRUE | 1.05 (1.00, 1.10) | 0.060 | FALSE | 1.18 (1.07, 1.30) | 6.64E-04 | FALSE | 1.37 (1.17, 1.59) | 5.61E-05 | TRUE | 1.79 (1.54, 2.08) | 3.61E-14 | TRUE | 1.43 (1.20, 1.71) | 6.58E-05 | TRUE |
| Myocardial infarction | 1.14 (1.07, 1.22) | 5.23E-05 | TRUE | 1.05 (0.97, 1.14) | 0.205 | FALSE | 1.30 (1.12, 1.50) | 4.54E-04 | FALSE | 1.65 (1.30, 2.09) | 3.41E-05 | TRUE | 2.31 (1.83, 2.91) | 2.27E-12 | TRUE | 1.75 (1.33, 2.30) | 6.10E-05 | TRUE |
| Pyogenic arthritis | 2.10 (1.41, 3.13) | 2.87E-04 | TRUE | 1.73 (1.07, 2.78) | 0.024 | FALSE | 2.66 (1.17, 6.08) | 2.00E-02 | FALSE | 1.71 (0.43, 6.80) | 0.444 | FALSE | 9.58 (2.44, 37.68) | 0.001 | FALSE | 6.13 (1.24, 30.22) | 0.026 | FALSE |
| Circulatory disease | 1.04 (1.02, 1.07) | 3.29E-04 | TRUE | 1.02 (0.99, 1.05) | 0.258 | FALSE | 1.10 (1.04, 1.16) | 6.94E-04 | FALSE | 1.31 (1.20, 1.43) | 1.59E-09 | TRUE | 1.29 (1.18, 1.41) | 9.47E-09 | TRUE | 1.17 (1.05, 1.29) | 0.003 | FALSE |
| Disorders of metabolism | 1.07 (1.03, 1.11) | 3.33E-04 | TRUE | 1.03 (0.99, 1.08) | 0.157 | FALSE | 1.17 (1.08, 1.27) | 1.06E-04 | TRUE | 1.12 (0.98, 1.27) | 0.100 | FALSE | 1.58 (1.39, 1.80) | 3.09E-12 | TRUE | 1.52 (1.30, 1.76) | 6.35E-08 | TRUE |
| Hypercholesterolemia | 1.08 (1.04, 1.12) | 3.34E-04 | TRUE | 1.00 (0.95, 1.05) | 0.913 | FALSE | 1.32 (1.20, 1.45) | 3.53E-09 | TRUE | 1.23 (1.06, 1.43) | 0.006 | FALSE | 1.90 (1.64, 2.20) | 8.73E-18 | TRUE | 1.84 (1.55, 2.19) | 3.01E-12 | TRUE |

**Supplementary Table S19. Sensitivity analysis by excluding the pleiotropic loci of metabolic traits.**

| **Disease outcomes** | **GRS of all urate loci (n=31)** | | | **GRS of loci without pleiotropy on obesity (n=21)** | | | **GRS of loci without pleiotropy on BP (n=21)** | | | **GRS of loci without pleiotropy on lipids (n=25)** | | | **GRS of loci without pleiotropy on glucose (n=28)** | | |
| --- | --- | --- | --- | --- | --- | --- | --- | --- | --- | --- | --- | --- | --- | --- | --- |
|  | **OR (95%CI)** | **p-value** | **FDR** | **OR (95%CI)** | **p-value** | **FDR** | **OR (95%CI)** | **p-value** | **FDR** | **OR (95%CI)** | **p-value** | **FDR** | **OR (95%CI)** | **p-value** | **FDR** |
| Gout | 5.37 (4.67, 6.18) | 4.27E-123 | TRUE | 3.89 (3.32, 4.56) | 1.01E-62 | TRUE | 5.42 (4.68, 6.28) | 6.00E-113 | TRUE | 5.07 (4.38, 5.86) | 5.27E-105 | TRUE | 5.09 (4.41, 5.88) | 1.31E-108 | TRUE |
| Inflammatory polyarthropathies | 1.27 (1.21, 1.34) | 4.97E-19 | TRUE | 1.21 (1.14, 1.28) | 6.43E-10 | TRUE | 1.26 (1.19, 1.33) | 1.02E-15 | TRUE | 1.26 (1.19, 1.33) | 1.16E-15 | TRUE | 1.27 (1.20, 1.34) | 1.34E-17 | TRUE |
| Hypertension | 1.07 (1.05, 1.11) | 6.02E-07 | TRUE | 1.07 (1.03, 1.10) | 2.31E-04 | FALSE | 1.04 (1.01, 1.07) | 2.24E-02 | FALSE | 1.05 (1.02, 1.09) | 0.002 | FALSE | 1.08 (1.05, 1.11) | 1.82E-06 | TRUE |
| Essential hypertension | 1.08 (1.05, 1.11) | 6.26E-07 | TRUE | 1.07 (1.03, 1.10) | 2.44E-04 | FALSE | 1.04 (1.01, 1.07) | 2.23E-02 | FALSE | 1.05 (1.02, 1.09) | 0.002 | FALSE | 1.08 (1.05, 1.11) | 1.83E-06 | TRUE |
| Coronary atherosclerosis | 1.10 (1.05, 1.14) | 1.17E-05 | TRUE | 1.08 (1.03, 1.13) | 0.001 | FALSE | 1.08 (1.03, 1.13) | 7.82E-04 | FALSE | 1.05 (1.01, 1.10) | 0.022 | FALSE | 1.08 (1.03, 1.13) | 4.88E-04 | FALSE |
| Gouty arthropathy | 5.10 (2.45, 10.66) | 1.39E-05 | TRUE | 4.04 (1.73, 9.44) | 0.001 | FALSE | 5.69 (2.61, 12.37) | 1.19E-05 | TRUE | 5.36 (2.47, 11.63) | 2.16E-05 | TRUE | 4.78 (2.24, 10.21) | 5.37E-05 | TRUE |
| Chronic Ischaemic heart disease | 1.09 (1.05, 1.14) | 1.52E-05 | TRUE | 1.08 (1.03, 1.13) | 0.002 | FALSE | 1.08 (1.03, 1.13) | 8.61E-04 | FALSE | 1.05 (1.01, 1.10) | 0.024 | FALSE | 1.08 (1.03, 1.13) | 5.58E-04 | FALSE |
| Ischaemic Heart Disease | 1.09 (1.05, 1.14) | 1.73E-05 | TRUE | 1.08 (1.03, 1.13) | 0.002 | FALSE | 1.08 (1.03, 1.13) | 9.51E-04 | FALSE | 1.05 (1.01, 1.10) | 0.026 | FALSE | 1.08 (1.03, 1.13) | 6.04E-04 | FALSE |
| Myocardial infarction | 1.14 (1.07, 1.22) | 5.23E-05 | TRUE | 1.11 (1.03, 1.20) | 0.006 | FALSE | 1.11 (1.04, 1.19) | 0.003 | FALSE | 1.08 (1.01, 1.16) | 0.033 | FALSE | 1.12 (1.04, 1.20) | 0.002 | FALSE |
| Pyogenic arthritis | 2.10 (1.41, 3.13) | 2.87E-04 | TRUE | 1.96 (1.24, 3.09) | 0.004 | FALSE | 2.14 (1.41, 3.26) | 3.73E-04 | FALSE | 1.82 (1.20, 2.77) | 0.005 | FALSE | 1.95 (1.29, 2.95) | 0.001 | FALSE |
| Circulatory disease | 1.04 (1.02, 1.07) | 3.29E-04 | TRUE | 1.03 (1.01, 1.06) | 0.020 | FALSE | 1.03 (1.00, 1.05) | 0.048 | FALSE | 1.03 (1.00, 1.05) | 0.041 | FALSE | 1.04 (1.01, 1.07) | 0.003 | FALSE |
| Disorders of metabolism | 1.07 (1.03, 1.11) | 3.33E-04 | TRUE | 1.04 (1.00, 1.09) | 0.038 | FALSE | 1.07 (1.03, 1.11) | 0.001 | FALSE | 1.03 (0.99, 1.07) | 0.094 | FALSE | 1.05 (1.01, 1.09) | 0.019 | FALSE |
| Hypercholesterolemia | 1.08 (1.04, 1.12) | 3.34E-04 | TRUE | 1.03 (0.98, 1.07) | 0.293 | FALSE | 1.07 (1.02, 1.12) | 0.003 | FALSE | 1.03 (0.98, 1.07) | 0.233 | FALSE | 1.04 (1.00, 1.09) | 0.051 | FALSE |

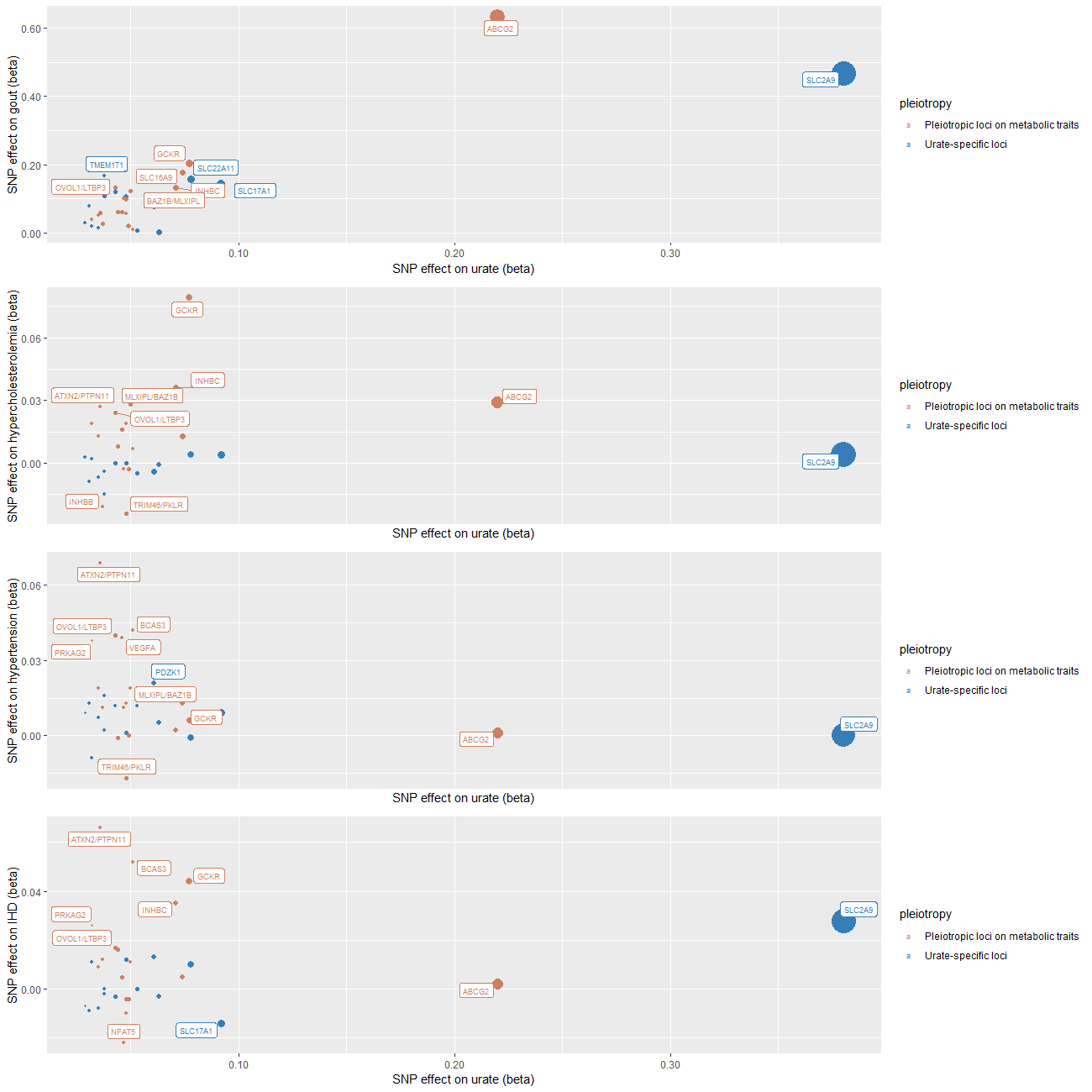

**Figure S1. Scatter plots of the effects of pleiotropic loci (mapped with genes) on serum urate levels against their effects on disease outcomes.** The size of the points was scaled to be inversely proportional to the standard errors of the effect sizes. Two urate transporter genes (SLC2A9 and ABCG2) are recognised as the leading loci driving the association with gout, the GCKR gene is the leading locus driving the association with hypercholesterolemia, and the PTPN11/ATXN2 gene is the leading locus driving the association with hypertension and ischaemic heart diseases.

5.	Kottgen A, Albrecht E, Teumer A, Vitart V, Krumsiek J, Hundertmark C, Pistis G, Ruggiero D, O'Seaghdha CM, Haller T, Yang Q, Tanaka T, Johnson AD, Kutalik Z, Smith AV, Shi J, Struchalin M, Middelberg RP, Brown MJ, Gaffo AL, Pirastu N, Li G, Hayward C, Zemunik T, Huffman J, Yengo L, Zhao JH, Demirkan A, Feitosa MF, Liu X, Malerba G, Lopez LM, van der Harst P, Li X, Kleber ME, Hicks AA, Nolte IM, Johansson A, Murgia F, Wild SH, Bakker SJ, Peden JF, Dehghan A, Steri M, Tenesa A, Lagou V, Salo P, Mangino M, Rose LM, Lehtimaki T, Woodward OM, Okada Y, Tin A, Muller C, Oldmeadow C, Putku M, Czamara D, Kraft P, Frogheri L, Thun GA, Grotevendt A, Gislason GK, Harris TB, Launer LJ, McArdle P, Shuldiner AR, Boerwinkle E, Coresh J, Schmidt H, Schallert M, Martin NG, Montgomery GW, Kubo M, Nakamura Y, Tanaka T, Munroe PB, Samani NJ, Jacobs DR, Jr., Liu K, D'Adamo P, Ulivi S, Rotter JI, Psaty BM, Vollenweider P, Waeber G, Campbell S, Devuyst O, Navarro P, Kolcic I, Hastie N, Balkau B, Froguel P, Esko T, Salumets A, Khaw KT, Langenberg C, Wareham NJ, Isaacs A, Kraja A, Zhang Q, Wild PS, Scott RJ, Holliday EG, Org E, Viigimaa M, Bandinelli S, Metter JE, Lupo A, Trabetti E, Sorice R, Doring A, Lattka E, Strauch K, Theis F, Waldenberger M, Wichmann HE, Davies G, Gow AJ, Bruinenberg M, Stolk RP, Kooner JS, Zhang W, Winkelmann BR, Boehm BO, Lucae S, Penninx BW, Smit JH, Curhan G, Mudgal P, Plenge RM, Portas L, Persico I, Kirin M, Wilson JF, Mateo Leach I, van Gilst WH, Goel A, Ongen H, Hofman A, Rivadeneira F, Uitterlinden AG, Imboden M, von Eckardstein A, Cucca F, Nagaraja R, Piras MG, Nauck M, Schurmann C, Budde K, Ernst F, Farrington SM, Theodoratou E, Prokopenko I, Stumvoll M, Jula A, Perola M, Salomaa V, Shin SY, Spector TD, Sala C, Ridker PM, Kahonen M, Viikari J, Hengstenberg C, Nelson CP, Meschia JF, Nalls MA, Sharma P, Singleton AB, Kamatani N, Zeller T, Burnier M, Attia J, Laan M, Klopp N, Hillege HL, Kloiber S, Choi H, Pirastu M, Tore S, Probst-Hensch NM, Volzke H, Gudnason V, Parsa A, Schmidt R, Whitfield JB, Fornage M, Gasparini P, Siscovick DS, Polasek O, Campbell H, Rudan I, Bouatia-Naji N, Metspalu A, Loos RJ, van Duijn CM, Borecki IB, Ferrucci L, Gambaro G, Deary IJ, Wolffenbuttel BH, Chambers JC, Marz W, Pramstaller PP, Snieder H, Gyllensten U, Wright AF, Navis G, Watkins H, Witteman JC, Sanna S, Schipf S, Dunlop MG, Tonjes A, Ripatti S, Soranzo N, Toniolo D, Chasman DI, Raitakari O, Kao WH, Ciullo M, Fox CS, Caulfield M, Bochud M, Gieger C. Genome-wide association analyses identify 18 new loci associated with serum urate concentrations. Nat Genet 2013;**45**(2):145-54.

6.	Willer CJ, Schmidt EM, Sengupta S, Peloso GM, Gustafsson S, Kanoni S, Ganna A, Chen J, Buchkovich ML, Mora S, Beckmann JS, Bragg-Gresham JL, Chang HY, Demirkan A, Den Hertog HM, Do R, Donnelly LA, Ehret GB, Esko T, Feitosa MF, Ferreira T, Fischer K, Fontanillas P, Fraser RM, Freitag DF, Gurdasani D, Heikkila K, Hypponen E, Isaacs A, Jackson AU, Johansson A, Johnson T, Kaakinen M, Kettunen J, Kleber ME, Li X, Luan J, Lyytikainen LP, Magnusson PKE, Mangino M, Mihailov E, Montasser ME, Muller-Nurasyid M, Nolte IM, O'Connell JR, Palmer CD, Perola M, Petersen AK, Sanna S, Saxena R, Service SK, Shah S, Shungin D, Sidore C, Song C, Strawbridge RJ, Surakka I, Tanaka T, Teslovich TM, Thorleifsson G, Van den Herik EG, Voight BF, Volcik KA, Waite LL, Wong A, Wu Y, Zhang W, Absher D, Asiki G, Barroso I, Been LF, Bolton JL, Bonnycastle LL, Brambilla P, Burnett MS, Cesana G, Dimitriou M, Doney ASF, Doring A, Elliott P, Epstein SE, Ingi Eyjolfsson G, Gigante B, Goodarzi MO, Grallert H, Gravito ML, Groves CJ, Hallmans G, Hartikainen AL, Hayward C, Hernandez D, Hicks AA, Holm H, Hung YJ, Illig T, Jones MR, Kaleebu P, Kastelein JJP, Khaw KT, Kim E, Klopp N, Komulainen P, Kumari M, Langenberg C, Lehtimaki T, Lin SY, Lindstrom J, Loos RJF, Mach F, McArdle WL, Meisinger C, Mitchell BD, Muller G, Nagaraja R, Narisu N, Nieminen TVM, Nsubuga RN, Olafsson I, Ong KK, Palotie A, Papamarkou T, Pomilla C, Pouta A, Rader DJ, Reilly MP, Ridker PM, Rivadeneira F, Rudan I, Ruokonen A, Samani N, Scharnagl H, Seeley J, Silander K, Stancakova A, Stirrups K, Swift AJ, Tiret L, Uitterlinden AG, van Pelt LJ, Vedantam S, Wainwright N, Wijmenga C, Wild SH, Willemsen G, Wilsgaard T, Wilson JF, Young EH, Zhao JH, Adair LS, Arveiler D, Assimes TL, Bandinelli S, Bennett F, Bochud M, Boehm BO, Boomsma DI, Borecki IB, Bornstein SR, Bovet P, Burnier M, Campbell H, Chakravarti A, Chambers JC, Chen YI, Collins FS, Cooper RS, Danesh J, Dedoussis G, de Faire U, Feranil AB, Ferrieres J, Ferrucci L, Freimer NB, Gieger C, Groop LC, Gudnason V, Gyllensten U, Hamsten A, Harris TB, Hingorani A, Hirschhorn JN, Hofman A, Hovingh GK, Hsiung CA, Humphries SE, Hunt SC, Hveem K, Iribarren C, Jarvelin MR, Jula A, Kahonen M, Kaprio J, Kesaniemi A, Kivimaki M, Kooner JS, Koudstaal PJ, Krauss RM, Kuh D, Kuusisto J, Kyvik KO, Laakso M, Lakka TA, Lind L, Lindgren CM, Martin NG, Marz W, McCarthy MI, McKenzie CA, Meneton P, Metspalu A, Moilanen L, Morris AD, Munroe PB, Njolstad I, Pedersen NL, Power C, Pramstaller PP, Price JF, Psaty BM, Quertermous T, Rauramaa R, Saleheen D, Salomaa V, Sanghera DK, Saramies J, Schwarz PEH, Sheu WH, Shuldiner AR, Siegbahn A, Spector TD, Stefansson K, Strachan DP, Tayo BO, Tremoli E, Tuomilehto J, Uusitupa M, van Duijn CM, Vollenweider P, Wallentin L, Wareham NJ, Whitfield JB, Wolffenbuttel BHR, Ordovas JM, Boerwinkle E, Palmer CNA, Thorsteinsdottir U, Chasman DI, Rotter JI, Franks PW, Ripatti S, Cupples LA, Sandhu MS, Rich SS, Boehnke M, Deloukas P, Kathiresan S, Mohlke KL, Ingelsson E, Abecasis GR, Global Lipids Genetics C. Discovery and refinement of loci associated with lipid levels. Nat Genet 2013;**45**(11):1274-1283.

7.	Shungin D, Winkler TW, Croteau-Chonka DC, Ferreira T, Locke AE, Magi R, Strawbridge RJ, Pers TH, Fischer K, Justice AE, Workalemahu T, Wu JMW, Buchkovich ML, Heard-Costa NL, Roman TS, Drong AW, Song C, Gustafsson S, Day FR, Esko T, Fall T, Kutalik Z, Luan J, Randall JC, Scherag A, Vedantam S, Wood AR, Chen J, Fehrmann R, Karjalainen J, Kahali B, Liu CT, Schmidt EM, Absher D, Amin N, Anderson D, Beekman M, Bragg-Gresham JL, Buyske S, Demirkan A, Ehret GB, Feitosa MF, Goel A, Jackson AU, Johnson T, Kleber ME, Kristiansson K, Mangino M, Leach IM, Medina-Gomez C, Palmer CD, Pasko D, Pechlivanis S, Peters MJ, Prokopenko I, Stancakova A, Sung YJ, Tanaka T, Teumer A, Van Vliet-Ostaptchouk JV, Yengo L, Zhang W, Albrecht E, Arnlov J, Arscott GM, Bandinelli S, Barrett A, Bellis C, Bennett AJ, Berne C, Bluher M, Bohringer S, Bonnet F, Bottcher Y, Bruinenberg M, Carba DB, Caspersen IH, Clarke R, Daw EW, Deelen J, Deelman E, Delgado G, Doney AS, Eklund N, Erdos MR, Estrada K, Eury E, Friedrich N, Garcia ME, Giedraitis V, Gigante B, Go AS, Golay A, Grallert H, Grammer TB, Grassler J, Grewal J, Groves CJ, Haller T, Hallmans G, Hartman CA, Hassinen M, Hayward C, Heikkila K, Herzig KH, Helmer Q, Hillege HL, Holmen O, Hunt SC, Isaacs A, Ittermann T, James AL, Johansson I, Juliusdottir T, Kalafati IP, Kinnunen L, Koenig W, Kooner IK, Kratzer W, Lamina C, Leander K, Lee NR, Lichtner P, Lind L, Lindstrom J, Lobbens S, Lorentzon M, Mach F, Magnusson PK, Mahajan A, McArdle WL, Menni C, Merger S, Mihailov E, Milani L, Mills R, Moayyeri A, Monda KL, Mooijaart SP, Muhleisen TW, Mulas A, Muller G, Muller-Nurasyid M, Nagaraja R, Nalls MA, Narisu N, Glorioso N, Nolte IM, Olden M, Rayner NW, Renstrom F, Ried JS, Robertson NR, Rose LM, Sanna S, Scharnagl H, Scholtens S, Sennblad B, Seufferlein T, Sitlani CM, Smith AV, Stirrups K, Stringham HM, Sundstrom J, Swertz MA, Swift AJ, Syvanen AC, Tayo BO, Thorand B, Thorleifsson G, Tomaschitz A, Troffa C, van Oort FV, Verweij N, Vonk JM, Waite LL, Wennauer R, Wilsgaard T, Wojczynski MK, Wong A, Zhang Q, Zhao JH, Brennan EP, Choi M, Eriksson P, Folkersen L, Franco-Cereceda A, Gharavi AG, Hedman AK, Hivert MF, Huang J, Kanoni S, Karpe F, Keildson S, Kiryluk K, Liang L, Lifton RP, Ma B, McKnight AJ, McPherson R, Metspalu A, Min JL, Moffatt MF, Montgomery GW, Murabito JM, Nicholson G, Nyholt DR, Olsson C, Perry JR, Reinmaa E, Salem RM, Sandholm N, Schadt EE, Scott RA, Stolk L, Vallejo EE, Westra HJ, Zondervan KT, Amouyel P, Arveiler D, Bakker SJ, Beilby J, Bergman RN, Blangero J, Brown MJ, Burnier M, Campbell H, Chakravarti A, Chines PS, Claudi-Boehm S, Collins FS, Crawford DC, Danesh J, de Faire U, de Geus EJ, Dorr M, Erbel R, Eriksson JG, Farrall M, Ferrannini E, Ferrieres J, Forouhi NG, Forrester T, Franco OH, Gansevoort RT, Gieger C, Gudnason V, Haiman CA, Harris TB, Hattersley AT, Heliovaara M, Hicks AA, Hingorani AD, Hoffmann W, Hofman A, Homuth G, Humphries SE, Hypponen E, Illig T, Jarvelin MR, Johansen B, Jousilahti P, Jula AM, Kaprio J, Kee F, Keinanen-Kiukaanniemi SM, Kooner JS, Kooperberg C, Kovacs P, Kraja AT, Kumari M, Kuulasmaa K, Kuusisto J, Lakka TA, Langenberg C, Le Marchand L, Lehtimaki T, Lyssenko V, Mannisto S, Marette A, Matise TC, McKenzie CA, McKnight B, Musk AW, Mohlenkamp S, Morris AD, Nelis M, Ohlsson C, Oldehinkel AJ, Ong KK, Palmer LJ, Penninx BW, Peters A, Pramstaller PP, Raitakari OT, Rankinen T, Rao DC, Rice TK, Ridker PM, Ritchie MD, Rudan I, Salomaa V, Samani NJ, Saramies J, Sarzynski MA, Schwarz PE, Shuldiner AR, Staessen JA, Steinthorsdottir V, Stolk RP, Strauch K, Tonjes A, Tremblay A, Tremoli E, Vohl MC, Volker U, Vollenweider P, Wilson JF, Witteman JC, Adair LS, Bochud M, Boehm BO, Bornstein SR, Bouchard C, Cauchi S, Caulfield MJ, Chambers JC, Chasman DI, Cooper RS, Dedoussis G, Ferrucci L, Froguel P, Grabe HJ, Hamsten A, Hui J, Hveem K, Jockel KH, Kivimaki M, Kuh D, Laakso M, Liu Y, Marz W, Munroe PB, Njolstad I, Oostra BA, Palmer CN, Pedersen NL, Perola M, Perusse L, Peters U, Power C, Quertermous T, Rauramaa R, Rivadeneira F, Saaristo TE, Saleheen D, Sinisalo J, Slagboom PE, Snieder H, Spector TD, Stefansson K, Stumvoll M, Tuomilehto J, Uitterlinden AG, Uusitupa M, van der Harst P, Veronesi G, Walker M, Wareham NJ, Watkins H, Wichmann HE, Abecasis GR, Assimes TL, Berndt SI, Boehnke M, Borecki IB, Deloukas P, Franke L, Frayling TM, Groop LC, Hunter DJ, Kaplan RC, O'Connell JR, Qi L, Schlessinger D, Strachan DP, Thorsteinsdottir U, van Duijn CM, Willer CJ, Visscher PM, Yang J, Hirschhorn JN, Zillikens MC, McCarthy MI, Speliotes EK, North KE, Fox CS, Barroso I, Franks PW, Ingelsson E, Heid IM, Loos RJ, Cupples LA, Morris AP, Lindgren CM, Mohlke KL. New genetic loci link adipose and insulin biology to body fat distribution. Nature 2015;**518**(7538):187-196.

8.	Manning AK, Hivert MF, Scott RA, Grimsby JL, Bouatia-Naji N, Chen H, Rybin D, Liu CT, Bielak LF, Prokopenko I, Amin N, Barnes D, Cadby G, Hottenga JJ, Ingelsson E, Jackson AU, Johnson T, Kanoni S, Ladenvall C, Lagou V, Lahti J, Lecoeur C, Liu Y, Martinez-Larrad MT, Montasser ME, Navarro P, Perry JR, Rasmussen-Torvik LJ, Salo P, Sattar N, Shungin D, Strawbridge RJ, Tanaka T, van Duijn CM, An P, de Andrade M, Andrews JS, Aspelund T, Atalay M, Aulchenko Y, Balkau B, Bandinelli S, Beckmann JS, Beilby JP, Bellis C, Bergman RN, Blangero J, Boban M, Boehnke M, Boerwinkle E, Bonnycastle LL, Boomsma DI, Borecki IB, Bottcher Y, Bouchard C, Brunner E, Budimir D, Campbell H, Carlson O, Chines PS, Clarke R, Collins FS, Corbaton-Anchuelo A, Couper D, de Faire U, Dedoussis GV, Deloukas P, Dimitriou M, Egan JM, Eiriksdottir G, Erdos MR, Eriksson JG, Eury E, Ferrucci L, Ford I, Forouhi NG, Fox CS, Franzosi MG, Franks PW, Frayling TM, Froguel P, Galan P, de Geus E, Gigante B, Glazer NL, Goel A, Groop L, Gudnason V, Hallmans G, Hamsten A, Hansson O, Harris TB, Hayward C, Heath S, Hercberg S, Hicks AA, Hingorani A, Hofman A, Hui J, Hung J, Jarvelin MR, Jhun MA, Johnson PC, Jukema JW, Jula A, Kao WH, Kaprio J, Kardia SL, Keinanen-Kiukaanniemi S, Kivimaki M, Kolcic I, Kovacs P, Kumari M, Kuusisto J, Kyvik KO, Laakso M, Lakka T, Lannfelt L, Lathrop GM, Launer LJ, Leander K, Li G, Lind L, Lindstrom J, Lobbens S, Loos RJ, Luan J, Lyssenko V, Magi R, Magnusson PK, Marmot M, Meneton P, Mohlke KL, Mooser V, Morken MA, Miljkovic I, Narisu N, O'Connell J, Ong KK, Oostra BA, Palmer LJ, Palotie A, Pankow JS, Peden JF, Pedersen NL, Pehlic M, Peltonen L, Penninx B, Pericic M, Perola M, Perusse L, Peyser PA, Polasek O, Pramstaller PP, Province MA, Raikkonen K, Rauramaa R, Rehnberg E, Rice K, Rotter JI, Rudan I, Ruokonen A, Saaristo T, Sabater-Lleal M, Salomaa V, Savage DB, Saxena R, Schwarz P, Seedorf U, Sennblad B, Serrano-Rios M, Shuldiner AR, Sijbrands EJ, Siscovick DS, Smit JH, Small KS, Smith NL, Smith AV, Stancakova A, Stirrups K, Stumvoll M, Sun YV, Swift AJ, Tonjes A, Tuomilehto J, Trompet S, Uitterlinden AG, Uusitupa M, Vikstrom M, Vitart V, Vohl MC, Voight BF, Vollenweider P, Waeber G, Waterworth DM, Watkins H, Wheeler E, Widen E, Wild SH, Willems SM, Willemsen G, Wilson JF, Witteman JC, Wright AF, Yaghootkar H, Zelenika D, Zemunik T, Zgaga L, Wareham NJ, McCarthy MI, Barroso I, Watanabe RM, Florez JC, Dupuis J, Meigs JB, Langenberg C. A genome-wide approach accounting for body mass index identifies genetic variants influencing fasting glycemic traits and insulin resistance. Nat Genet 2012;**44**(6):659-69.

9.	Ehret GB, Munroe PB, Rice KM, Bochud M, Johnson AD, Chasman DI, Smith AV, Tobin MD, Verwoert GC, Hwang SJ, Pihur V, Vollenweider P, O'Reilly PF, Amin N, Bragg-Gresham JL, Teumer A, Glazer NL, Launer L, Zhao JH, Aulchenko Y, Heath S, Sober S, Parsa A, Luan J, Arora P, Dehghan A, Zhang F, Lucas G, Hicks AA, Jackson AU, Peden JF, Tanaka T, Wild SH, Rudan I, Igl W, Milaneschi Y, Parker AN, Fava C, Chambers JC, Fox ER, Kumari M, Go MJ, van der Harst P, Kao WH, Sjogren M, Vinay DG, Alexander M, Tabara Y, Shaw-Hawkins S, Whincup PH, Liu Y, Shi G, Kuusisto J, Tayo B, Seielstad M, Sim X, Nguyen KD, Lehtimaki T, Matullo G, Wu Y, Gaunt TR, Onland-Moret NC, Cooper MN, Platou CG, Org E, Hardy R, Dahgam S, Palmen J, Vitart V, Braund PS, Kuznetsova T, Uiterwaal CS, Adeyemo A, Palmas W, Campbell H, Ludwig B, Tomaszewski M, Tzoulaki I, Palmer ND, Aspelund T, Garcia M, Chang YP, O'Connell JR, Steinle NI, Grobbee DE, Arking DE, Kardia SL, Morrison AC, Hernandez D, Najjar S, McArdle WL, Hadley D, Brown MJ, Connell JM, Hingorani AD, Day IN, Lawlor DA, Beilby JP, Lawrence RW, Clarke R, Hopewell JC, Ongen H, Dreisbach AW, Li Y, Young JH, Bis JC, Kahonen M, Viikari J, Adair LS, Lee NR, Chen MH, Olden M, Pattaro C, Bolton JA, Kottgen A, Bergmann S, Mooser V, Chaturvedi N, Frayling TM, Islam M, Jafar TH, Erdmann J, Kulkarni SR, Bornstein SR, Grassler J, Groop L, Voight BF, Kettunen J, Howard P, Taylor A, Guarrera S, Ricceri F, Emilsson V, Plump A, Barroso I, Khaw KT, Weder AB, Hunt SC, Sun YV, Bergman RN, Collins FS, Bonnycastle LL, Scott LJ, Stringham HM, Peltonen L, Perola M, Vartiainen E, Brand SM, Staessen JA, Wang TJ, Burton PR, Soler Artigas M, Dong Y, Snieder H, Wang X, Zhu H, Lohman KK, Rudock ME, Heckbert SR, Smith NL, Wiggins KL, Doumatey A, Shriner D, Veldre G, Viigimaa M, Kinra S, Prabhakaran D, Tripathy V, Langefeld CD, Rosengren A, Thelle DS, Corsi AM, Singleton A, Forrester T, Hilton G, McKenzie CA, Salako T, Iwai N, Kita Y, Ogihara T, Ohkubo T, Okamura T, Ueshima H, Umemura S, Eyheramendy S, Meitinger T, Wichmann HE, Cho YS, Kim HL, Lee JY, Scott J, Sehmi JS, Zhang W, Hedblad B, Nilsson P, Smith GD, Wong A, Narisu N, Stancakova A, Raffel LJ, Yao J, Kathiresan S, O'Donnell CJ, Schwartz SM, Ikram MA, Longstreth WT, Jr., Mosley TH, Seshadri S, Shrine NR, Wain LV, Morken MA, Swift AJ, Laitinen J, Prokopenko I, Zitting P, Cooper JA, Humphries SE, Danesh J, Rasheed A, Goel A, Hamsten A, Watkins H, Bakker SJ, van Gilst WH, Janipalli CS, Mani KR, Yajnik CS, Hofman A, Mattace-Raso FU, Oostra BA, Demirkan A, Isaacs A, Rivadeneira F, Lakatta EG, Orru M, Scuteri A, Ala-Korpela M, Kangas AJ, Lyytikainen LP, Soininen P, Tukiainen T, Wurtz P, Ong RT, Dorr M, Kroemer HK, Volker U, Volzke H, Galan P, Hercberg S, Lathrop M, Zelenika D, Deloukas P, Mangino M, Spector TD, Zhai G, Meschia JF, Nalls MA, Sharma P, Terzic J, Kumar MV, Denniff M, Zukowska-Szczechowska E, Wagenknecht LE, Fowkes FG, Charchar FJ, Schwarz PE, Hayward C, Guo X, Rotimi C, Bots ML, Brand E, Samani NJ, Polasek O, Talmud PJ, Nyberg F, Kuh D, Laan M, Hveem K, Palmer LJ, van der Schouw YT, Casas JP, Mohlke KL, Vineis P, Raitakari O, Ganesh SK, Wong TY, Tai ES, Cooper RS, Laakso M, Rao DC, Harris TB, Morris RW, Dominiczak AF, Kivimaki M, Marmot MG, Miki T, Saleheen D, Chandak GR, Coresh J, Navis G, Salomaa V, Han BG, Zhu X, Kooner JS, Melander O, Ridker PM, Bandinelli S, Gyllensten UB, Wright AF, Wilson JF, Ferrucci L, Farrall M, Tuomilehto J, Pramstaller PP, Elosua R, Soranzo N, Sijbrands EJ, Altshuler D, Loos RJ, Shuldiner AR, Gieger C, Meneton P, Uitterlinden AG, Wareham NJ, Gudnason V, Rotter JI, Rettig R, Uda M, Strachan DP, Witteman JC, Hartikainen AL, Beckmann JS, Boerwinkle E, Vasan RS, Boehnke M, Larson MG, Jarvelin MR, Psaty BM, Abecasis GR, Chakravarti A, Elliott P, van Duijn CM, Newton-Cheh C, Levy D, Caulfield MJ, Johnson T. Genetic variants in novel pathways influence blood pressure and cardiovascular disease risk. Nature 2011;**478**(7367):103-9.

10.	Malik R, Traylor M, Pulit SL, Bevan S, Hopewell JC, Holliday EG, Zhao W, Abrantes P, Amouyel P, Attia JR, Battey TW, Berger K, Boncoraglio GB, Chauhan G, Cheng YC, Chen WM, Clarke R, Cotlarciuc I, Debette S, Falcone GJ, Ferro JM, Gamble DM, Ilinca A, Kittner SJ, Kourkoulis CE, Lemmens R, Levi CR, Lichtner P, Lindgren A, Liu J, Meschia JF, Mitchell BD, Oliveira SA, Pera J, Reiner AP, Rothwell PM, Sharma P, Slowik A, Sudlow CL, Tatlisumak T, Thijs V, Vicente AM, Woo D, Seshadri S, Saleheen D, Rosand J, Markus HS, Worrall BB, Dichgans M. Low-frequency and common genetic variation in ischemic stroke: The METASTROKE collaboration. Neurology 2016;**86**(13):1217-26.

11.	Hemani G, Bowden J, Haycock P, Zheng J, Davis O, Flach P, Gaunt T, Smith GD. Automating Mendelian randomization through machine learning to construct a putative causal map of the human phenome. bioRxiv 2017.

12.	Bowden J, Davey Smith G, Burgess S. Mendelian randomization with invalid instruments: effect estimation and bias detection through Egger regression. Int J Epidemiol 2015;**44**(2):512-25.

13.	Bowden J, Del Greco MF, Minelli C, Davey Smith G, Sheehan N, Thompson J. A framework for the investigation of pleiotropy in two-sample summary data Mendelian randomization. Stat Med 2017;**36**(11):1783-1802.

14.	Bowden J, Davey Smith G, Haycock PC, Burgess S. Consistent Estimation in Mendelian Randomization with Some Invalid Instruments Using a Weighted Median Estimator. Genet Epidemiol 2016;**40**(4):304-14.

15.	Hartwig FP, Davey Smith G, Bowden J. Robust inference in summary data Mendelian randomization via the zero modal pleiotropy assumption. Int J Epidemiol 2017;**46**(6):1985-1998.
